# Frugivore gut passage increases seed germination: an updated meta-analysis

**DOI:** 10.1101/2021.10.12.462022

**Authors:** Haldre S. Rogers, Brittany R. Cavazos, Ann Marie Gawel, Alex Karnish, Courtenay A. Ray, Ethan Rose, Hugo Thierry, Evan C. Fricke

## Abstract

Many plants rely on animal mutualists for reproduction. Quantifying how animal mutualists impact plant performance provides a foundation for modelling how change in animal communities affects the composition and functioning of plant communities. We performed a meta-analysis of 2539 experiments, 6 times more than the last comprehensive meta-analysis, examining how gut passage by frugivores influences seed germination. We simultaneously analyzed multiple predictor variables related to study methodology, location, and frugivore identity to disentangle methodological from ecological impacts on effect sizes. We found that gut passage by birds, fish, reptiles, bats, primates, and other mammals on average increased seed germination, but that the magnitude varied across vertebrate groups. The positive effects of gut passage were largely explained by the de-inhibitory effects of pulp removal rather than by the scarification of seed tissues. Some previous studies and meta-analyses that found no effect of gut passage only tested scarification or did not distinguish between these tests of scarification and pulp removal. We found that, for a typical fleshy-fruited plant species, the lack of gut passage reduces germination by 60%. From an evolutionary perspective, this indicates a large risk associated with reliance on animal mutualists that is balanced against the benefits of animal-mediated seed dispersal. From a conservation perspective, this highlights the potential for large demographic consequences of frugivore declines on plant populations. Our database and findings advance quantitative predictions for the role of fruit-frugivore interactions in shaping plant communities in the Anthropocene.

## INTRODUCTION

Interactions among species support the functioning of ecosystems (Loreau *et al.* 2001). Mutualisms like pollination and seed dispersal affect plant communities by impacting both plant reproduction and the movement of genetic material (Kremen *et al.* 2007). Mutualists thus shape the ecosystem services that plants provide to people (Garibaldi *et al.* 2013; Egerer *et al.* 2017). Mutualist declines under anthropogenic change threaten plant diversity, ecosystem functioning, and ecosystem services (Potts *et al.* 2010; Rogers *et al.* 2021). Generalizing knowledge of the functional impacts of such species interactions informs how mutualistic interactions affect community dynamics and the consequences of ongoing declines of animal mutualists (Brodie *et al.* 2018). The outcome of a species interaction for ecosystem function depends on the frequency of interaction (quantity component) as well as the net impact of that interaction on individual plant performance (quality component), which could be positive, neutral, or negative even for putatively ‘mutualistic’ interactions. Formalized in the effectiveness framework (Schupp *et al.* 2017), the product of the quantity and quality components gives an estimate of the total impact of the interactions among a species pair on ecosystem function. Whereas the quantity component may be more easily measured (i.e., via direct observation), measuring the quality component requires intensive experiments that track impacts on reproduction or survival over months or years.

Seed dispersal by animals is widespread across the phylogeny of seed plants (Jordano 2000; Rogers *et al.* 2021). Roughly half of the ~350,000 angiosperm species producing fleshy fruit (Aslan *et al.* 2013) are adapted for consumption and dispersal by mutualistic partners including birds, non-avian reptiles, bats, primates, and invertebrates. Global change factors including overhunting, species invasion, and fragmentation are causing declines in many seed-dispersing mutualists (McConkey *et al.* 2012), with particularly striking declines in large-bodied mammals and birds (McConkey *et al.* 2012; Galetti & Dirzo 2013; Dirzo *et al.* 2014). Changes in frugivore abundance and diversity may affect plant populations through mechanisms such as reduced seedling recruitment due to insufficient seed consumption and gut passage, reduced movement away from areas of high mortality near conspecific parents or to areas suitable for germination, and reduced colonization ability (Farwig & Berens 2012; Aslan *et al.* 2019; Rogers *et al.* 2021).

The impact of frugivore gut passage on seed germination has been studied for more than a century (Barrows & Schwarz 1895 p. 85-87; Troup 1921; Ridley 1930; Krefting & Roe 1949). Experiments measuring these impacts compare the probability of germination of gut-passed seeds to that of seeds that are not gut-passed (Samuels & Levey 2005). These experiments address several basic ecological and evolutionary questions. First, they address some of the fundamental costs and benefits to plants that result from engaging in mutualistic seed dispersal interactions, specifically the cost of seed destruction and benefit of increased seed germination by frugivores. Second, they examine aspects of seed and reproductive biology including the impact on the probability or timing of germination due to removal of inhibitory cues through pulp removal (de-inhibition effect) versus that of mechanical and chemical changes to seed tissues (scarification effect). Third, they can reveal how phylogenetically or morphologically distinct animal partners vary in their impacts on germination, offering insights into coevolutionary processes among mutualists. Further, these experiments also have direct applications in conservation contexts. Studies on individual plant or animal species elucidate the demographic consequences of frugivore declines; plant species more heavily dependent on frugivore gut passage are more vulnerable to mutualism disruption (Rogers *et al.* 2021) and frugivores that provide the largest functional benefits are of particular importance in conservation or restoration settings (Samuels & Levey 2005).

Previous reviews and meta-analyses have addressed the impact of gut passage on germination, covering studies across all frugivores (Traveset 1998 [315 experiments from 80 studies]; Traveset & Verdu 2002 [351 experiments from 83 studies]; Verdu & Traveset 2004 [216 experiments]; Soltani *et al.* 2018 [581 experiments from 76 studies]) and particular taxonomic groups of frugivores (primates - Fuzessy *et al.* 2016 [460 experiments from 19 studies]; bats - Saldaña-Vázquez *et al.* 2019 [106 experiments from 33 studies]). In general, these studies support a positive effect of gut passage for most plant species, with variation between frugivore groups. However, bats and reptiles were poorly represented in the last comprehensive meta-analysis (Traveset & Verdu 2002 [bats - 19 studies, reptiles - 39 studies]) and fish and insects were not included due to a lack of studies. A recent meta-analysis on bats (Saldaña-Vázquez *et al.* 2019) covered 5 times more experiments than Traveset and Verdu (2002) and concluded that gut passage by bats had a neutral effect on germination, which differs from the positive effect found by Traveset and Verdu (2002). In addition to the large number of studies published in the years since the last meta-analysis, updated analytic approaches (Viechtbauer 2010) allow for more robust insights. In particular, previous meta-analyses assess a single predictor variable at a time, whereas the metafor package (Viechtbauer 2010) facilitates the incorporation of multiple predictor variables in a single model.

A fundamental methodological limitation of prior meta-analyses, and of the vast majority of experiments included in the meta-analyses, is the use of manually de-pulped seeds for the control treatment, or even more problematically, the lack of distinction between manually de-pulped seeds and seeds within whole fruit. An experimental design that only uses manually de-pulped seeds for comparison with gut-passed poorly represents the ecosystem functioning provided by frugivores—or consequences of mutualist loss—because animals are responsible for both pulp removal and scarification in nature (Samuels & Levey 2005; Costa-Pereira 2017). Comparisons between gut-passed and manually de-pulped seeds, without comparison to whole fruit, fail to quantify the de-inhibitory effects of frugivores and may lead to incomplete conclusions on the impacts of gut passage. This is likely to lead to inaccurate conclusions if studies on specific animal groups disproportionately employ methods focused only on scarification or de-inhibition, because the confounded effects of study design and animal group would obscure differences between groups. The recent meta-analysis of studies involving bats was only able to include comparisons between gut-passed and manually de-pulped seeds, due to limited studies involving comparisons to whole fruits, and showed no overall effect of gut passage on germination (Saldaña-Vázquez *et al.* 2019). However, this only tested the scarification component of gut passage so it is premature to conclude that gut passage by bats does not affect germination.

Here, we compiled a database of all available studies testing the effect of gut passage on germination of fleshy-fruited plant species. The database includes 2539 experimental comparisons from 339 publications, an increase of 2188 experiments and 256 studies since the last comprehensive meta-analysis. By conducting the first meta-analysis to simultaneously analyze multiple predictor variables related to study methodology, location, and frugivore identity, we disentangle methodological from ecological impacts on effect sizes. This allows us to 1) compare the magnitude of the de-inhibition and scarification effects, 2) understand the effects of different frugivore taxa on mutualistic ecosystem functioning, and 3) examine coarse macroecological variation in effect sizes.

## METHODS

### Database compilation

We aimed to compile all primary experiments published through the end of 2017 on the impacts of animal gut passage on germination. To identify potential papers for inclusion in the meta-analysis, we performed a SCOPUS search using the search terms: “TITLE-ABS-KEY (germinat* AND (“seed dispers*” OR frugivor* OR “gut pass*” OR “ingest*” OR “endozoochor*”)))”. We supplemented this with studies cited in, or that cited, Traveset (1998). Among the 2,410 potential papers, we selected studies that compared germination of ingested seeds against a control, either whole fruit or manually de-pulped seeds for inclusion in this paper. We included studies where seeds were regurgitated following ingestion in addition to the great majority of cases where seeds were defecated. We analyzed data for studies where the proportion of seeds germinating could be discerned, such as the number of seeds sown and germinated in the gut-passed and control treatments or percent of seeds germinated. When data were only presented in figures, we used WebPlotDigitizer (Rohatgi 2017) to obtain quantitative values. We include a list of the 339 studies used in our meta-analysis in the Supplemental Materials (Table S1).

Along with data to characterize effect sizes from each study, we recorded several other variables related to the study or focal species. For methodological variables, we recorded the control that gut-passed seeds were compared to (whole fruit or mechanically cleaned seeds). Other less common treatments or experimental setups (e.g., comparisons to chemically scarified seeds) were excluded from analysis. We noted whether gut-passed seeds were collected by searching for scat in the field (field-collected) or were collected during feeding trials with captive animals (captive trial samples). We recorded the medium in which seeds were sown, either petri dishes, greenhouse soil, field soil, or other planting mediums (such as tree branches for mistletoe seeds). We recorded the plant and animal names to the finest taxonomic resolution available down to the species level, resolving taxonomy using the Taxonstand package in R (Cayuela *et al.* 2012). We assigned animal species to several animal groups: bird, reptile, bat, primate, other mammal, fish, and invertebrates. Using the Global Invasive Species Database, we determined whether each plant or animal is known to occur as an invasive species in any part of its current range. We noted the latitudinal region in which the study occurred (tropical, subtropical, temperate) and whether it occurred on an island or mainland ecosystem. We sought to understand how the number of animal species studied relates to the total number of frugivorous species. Focusing on birds and mammals, we recorded the IUCN Red List status of each studied animal species and for all bird and mammal species that have fruit in their diet (>5%) based on the EltonTraits 1.0 database (Wilman *et al.* 2014).

### Meta-analysis methods

We fit meta-analytic multivariate mixed effects models using the ‘rma.mv’ function in the metafor package in R (Viechtbauer 2010). The effect sizes were calculated as an odds ratio based on the number of seeds and germinants in the gut-passed and control treatments. When only the proportion germinating—not the absolute number—was reported, we assumed that the number of seeds in the experiment equaled the median number of seeds across experiments where these data were reported. Although this decision could influence sampling variances estimated for each experiment, this decision likely did not affect our conclusions because models run after excluding these cases gave qualitatively equivalent results. In a full model where we allowed random intercepts by plant and animal species, we included fixed effects describing the control type, feeding trial type, sowing medium, frugivore taxon, invasive status, latitude region, and mainland vs. island study location. To develop a best-fit model, we compared all nested models with fewer fixed effects and removed variables that did not improve AIC by 2 units. To evaluate the potential for phylogenetic non-independence to bias our conclusion, we ran equivalent models with a variance-covariance matrix based on the plant phylogeny. We constructed the dated plant phylogenetic tree using Phylomatic (Webb & Donoghue 2005) and the bladJ algorithm (Webb *et al.* 2008).

Using the best-fit model, we made specific comparisons outlined in the introduction using linear hypothesis testing in the ‘multcomp’ package in R (Hothorn *et al.* 2008). We assessed differences between the de-inhibition effect (difference between whole fruit and manually de-pulped germination) and the scarification effect (difference between de-pulped and gut-passed germination). We likewise tested for differences across the methodological factors (e.g., whether effect sizes differed among each combination of planting mediums), species-level factors (e.g., whether effect sizes differed among each pair of animal groups), and variables related to study location. For visualization of linear hypothesis test results, we obtained model estimates for a combination of levels of the categorical variables that characterize the total gut passage effect (de-inhibition and scarification) for a typical experiment. Specifically, this combination of levels represents a trial involving birds in the tropics, comparing to a whole fruit control, using captive feeding trials, and with seeds sown in petri dishes. Other combinations of levels would give identical statistical results for the linear comparisons because we did not allow interaction terms in the meta-analytic model.

We used two approaches to assess potential publication bias. We present histograms of the log odds effect size across all combinations and this effect size weighted by the inverse of the variance. These can indicate publication bias against studies with small effect sizes if depressed near zero. We also present a funnel plot, which can indicate publication bias if asymmetric. As a statistical test of funnel plot asymmetry, we present a rank correlation test (Begg and Mazumdar 1994).

## RESULTS

We analyzed data from 339 publications reporting the results of 2539 experimental comparisons between gut-passed and control seeds involving 1622 unique plant-frugivore interactions from 446 plant genera and 226 animal genera. The countries in which experiments were performed exhibit spatial heterogeneity, with the most well-studied countries including Brazil, the United States of America, Spain, Australia, and South Africa (Fig. 1a). The number of experiments per year has increased over time (Fig. 1b). Out of the total number of frugivorous bird and mammal species, the portion that has been the focus of a gut passage experiment is small (Fig. 1c). Whereas mammals have been studied roughly in proportion to IUCN Red List status, birds that are more threatened are disproportionately poorly studied (Fig. 1c).

**Figure 1.**
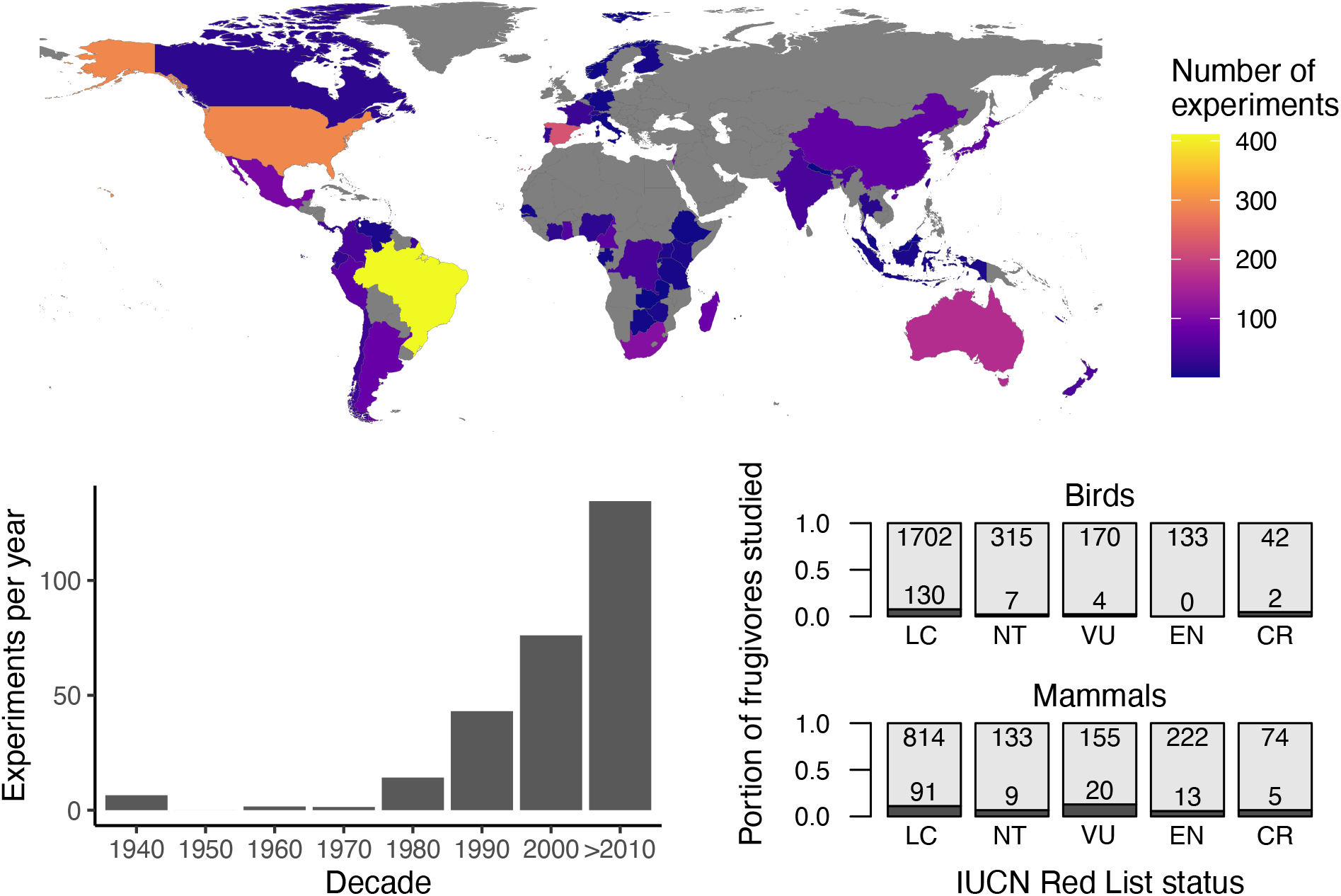
Study intensity in relation to space, time, and IUCN Red List status of frugivores. a) The number of experiments per country is shown on a color gradient; countries with no studies in the meta-analysis are shown in grey. b) The number of experiments per year has increased since the 1980s. c) The portion of bird and mammal species with fruit in their diet that have been assessed in a gut passage germination experiment (shown in dark grey; species count shown by numbers at bottom of each bar) is small relative to the total number of frugivorous species (shown in light grey; species counts at top of bar) and varies across IUCN Red List status.

We fit a meta-analytic mixed effects model using all predictor variables related to study methods, frugivore taxon, plant and animal invasive species status, and study location (Fig. S1, Table S1). An equivalent model with a covariance matrix based on plant phylogeny showed qualitatively and quantitatively similar results (Fig. S2, Table S2), suggesting that plant phylogenetic non-independence is unlikely to bias our conclusions. The one difference was a relative inflation of confidence intervals around the model intercept in the phylogenetic model. This may be due to closely related species, or the same species, exhibiting variable effect sizes in different experiments. The best fit model included all predictor variables except the variables describing whether the plant or animal species was an invasive species; neither variable predicted the effect of gut passage on germination (Fig. S1). We used this best fit model for linear hypothesis tests. The funnel plot did not suggest bias against publications with small effect sizes (Fig. S1d), but we did find evidence for funnel plot asymmetry (Kendall’s tau = 0.1025, p < 0.001).

Aspects of study design altered the measured impact of gut passage on germination (Fig. 2, Table S3). Effect sizes varied with the planting medium in which experimental seeds were sown, with seeds sown in petri dishes and other locations (e.g., tree branches for mistletoe seeds) showing more positive effect sizes than seeds sown in nursery or field soil (Fig. 2a). There was no significant difference in gut passage effects between studies where gut-passed seeds were collected during feeding trials with captive animals and where gut-passed seeds were collected from scat in the field (Fig. 2b).

**Figure 2.**
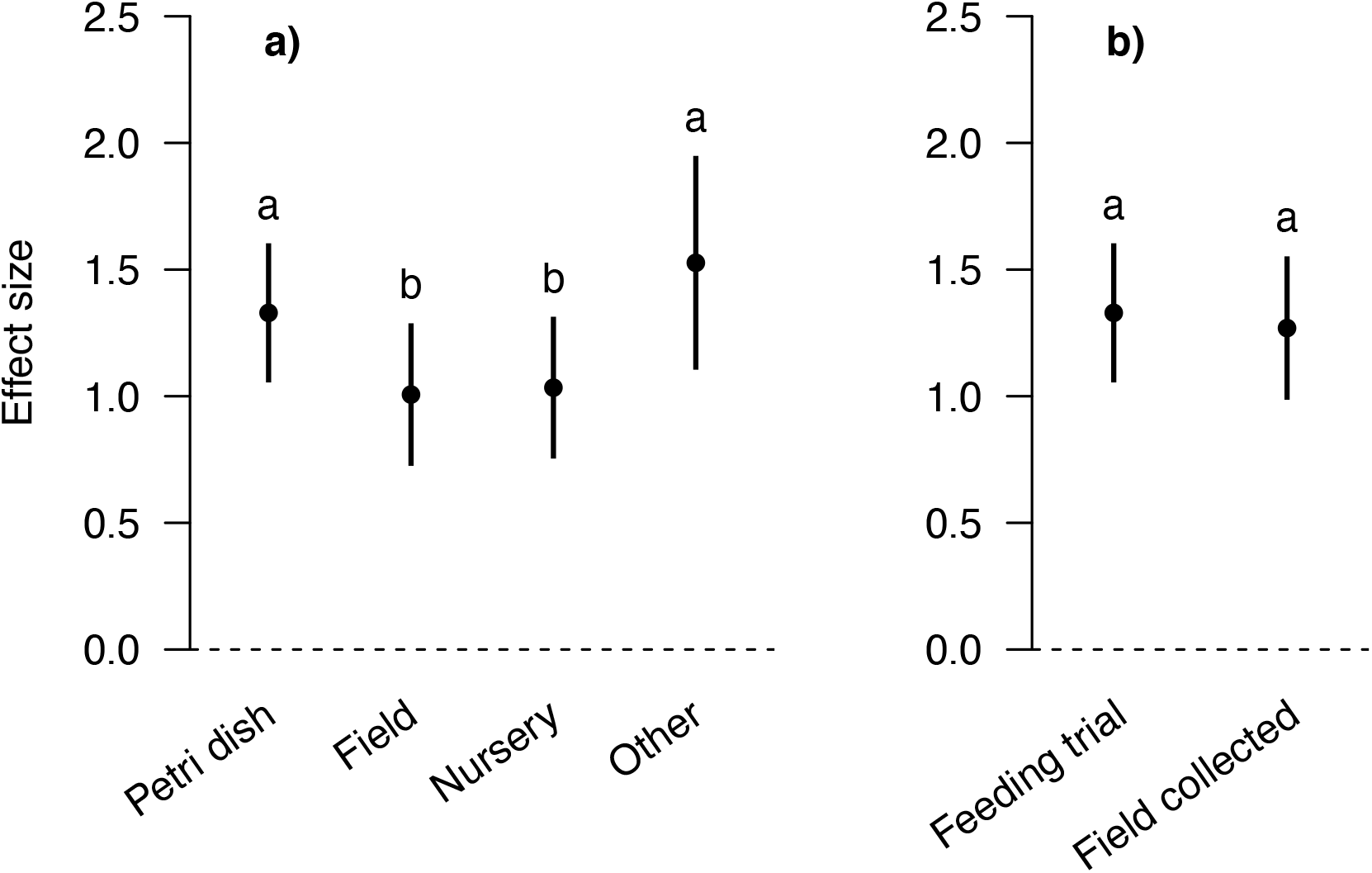
Impacts of study methodology on estimated effect sizes. Points represent model-estimated impacts of gut passage on germination (log-odds scale), bars indicate confidence intervals, and letters show statistically significant differences. Estimates were developed using values for the reference predictor combination and varying either (a) the planting medium or (b) whether the test used seeds from a feeding trial or field-collected seeds.

We modeled the scarification effect as the difference between the germination of gut-passed seeds versus seeds that were manually de-pulped and the de-inhibition effect as the difference between the germination of seeds in whole fruit versus manually de-pulped seeds. The scarification effect was significantly smaller than the de-inhibition effect (Fig. 3, Table S3).

**Figure 3.**
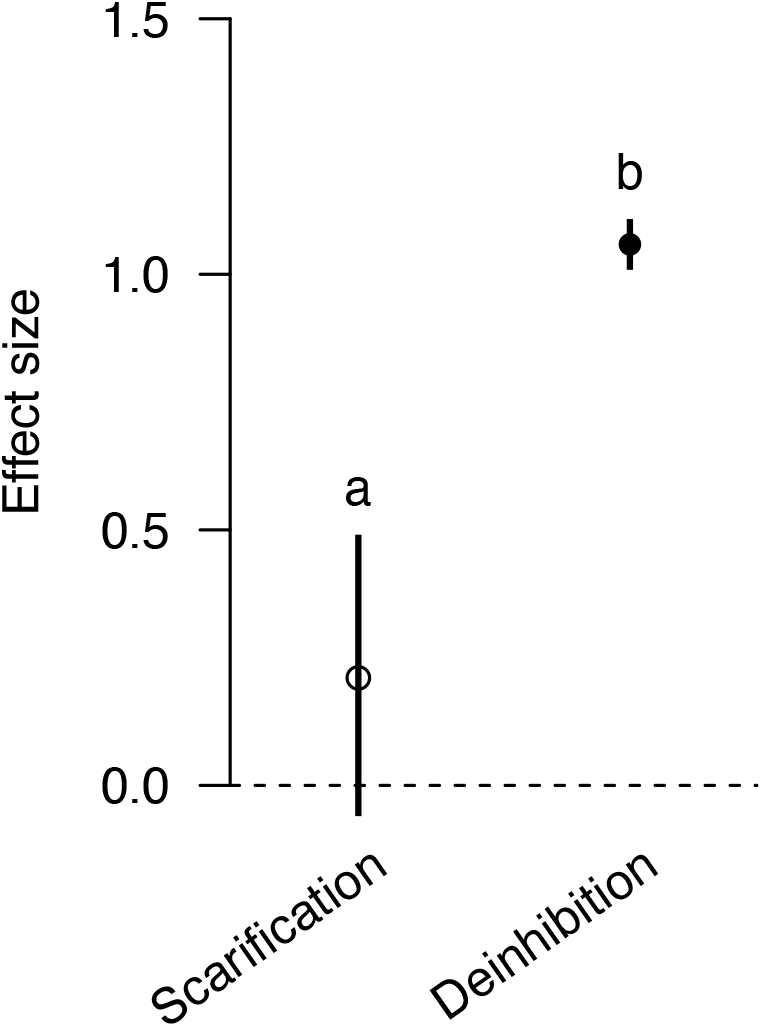
Disentangling the scarification and de-inhibition effects. Effects of scarification (caused by gut passage) were smaller than de-inhibition effects caused by pulp removal (independent of scarification via gut passage).

Other differences in effect sizes were explained by the frugivore group and study location (Fig. 4, Table S3). When considering the comprehensive effect of gut passage (both de-inhibition and scarification effects) on germination, birds exhibited positive effect sizes (Fig. 4a). Primates showed more positive effect sizes than birds, and bat effect sizes were similar to those of both birds and primates. Other mammals exhibited smaller but positive effect sizes. Reptiles had effect sizes similar to those of birds and bats. The positive effect sizes of fish could not be distinguished from the effect sizes of other vertebrate taxa. Invertebrates had negative mean effects exhibiting a marginally significant difference from zero. Latitudinal zone impacted effect sizes, with subtropical and temperate effect sizes similar to each other but both larger than tropical effect sizes (Fig. 4b). Mainland effect sizes were more positive than island effect sizes (Fig. 4c).

**Figure 4.**
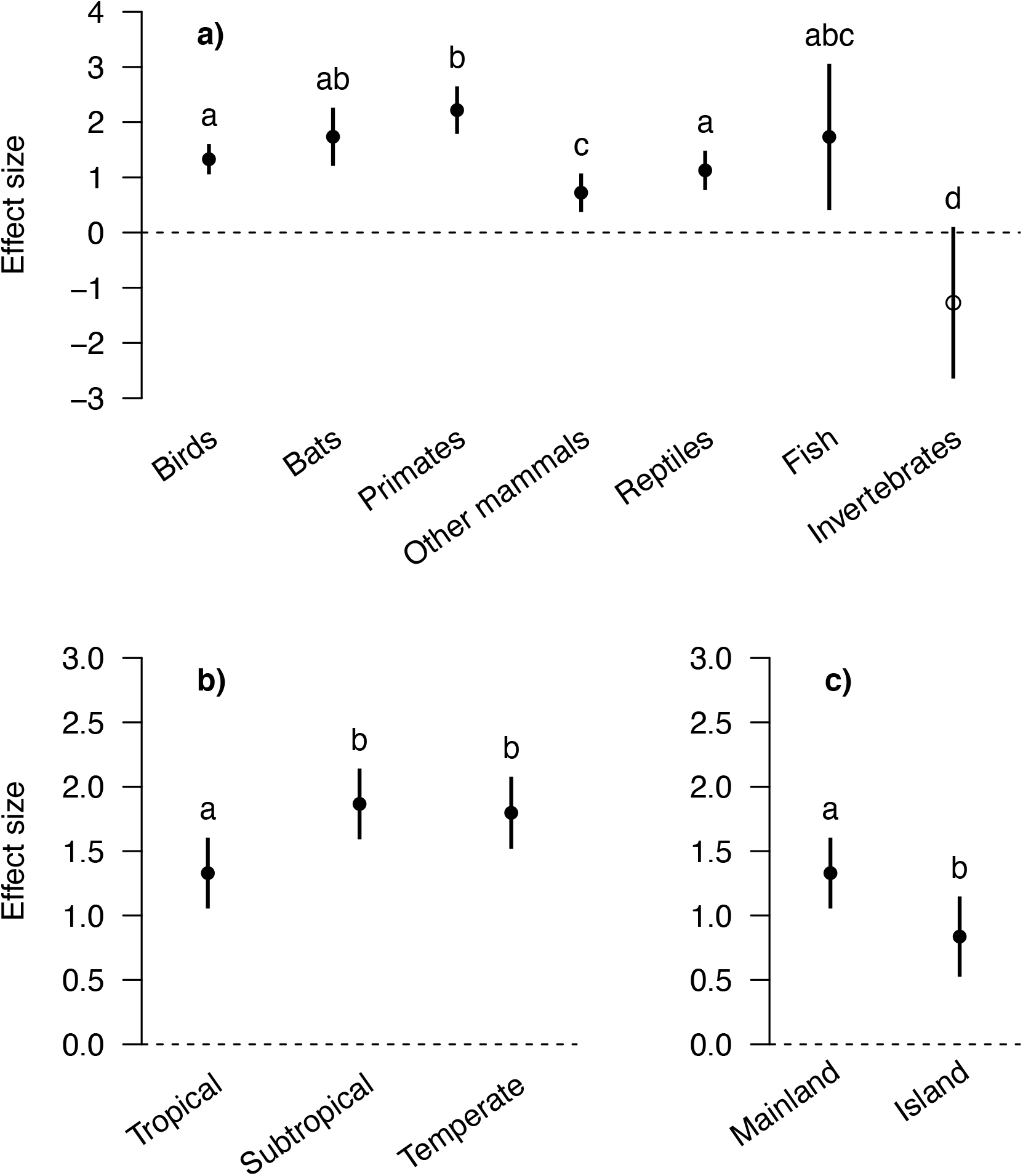
Differences in effect sizes explained by (a) frugivore taxon and (b, c) location.

## DISCUSSION

Fruit-frugivore interactions are widespread and increasingly affected by global change factors such as overhunting, invasive species, and habitat fragmentation (McConkey *et al.* 2012; Fricke & Svenning 2020; Rogers *et al.* 2021); however, the quantitative effect of these changes in animal communities on plant ecosystem functioning is poorly known. To quantify how frugivores influence plant performance through impacts of gut passage on germination, we performed a meta-analysis of more than 2500 experiments comparing germination of gut-passed seeds to seeds that were not passed by frugivores. Using the first meta-analytic approach to take multiple explanatory variables into account simultaneously, we found that frugivorous vertebrates exhibited on average strong positive effects of gut passage on germination whereas the few experiments with invertebrates showed a negative effect. Since about half of all plant species have fleshy fruit adapted for animal dispersal largely by vertebrates, defaunation of vertebrate frugivores will cascade to affect plant germination and recruitment.

Our meta-analysis is the first to separate the effects of de-inhibition through pulp removal from those of scarification from gut passage. Studies typically compare germination of gut-passed seeds to two types of ‘control’ treatments: either seeds that were mechanically cleaned of pulp by researchers or seeds that were left within whole fruits. The former quantifies the scarification effect—caused by physical or chemical changes to seed tissues—whereas the latter quantifies the sum of the scarification effect and de-inhibition effect—caused by removal of pulp and chemical cues within it (Samuels & Levey 2005; Robertson *et al.* 2006). Ultimately, frugivores both remove pulp and scarify the seed, so the most ecologically relevant comparison for testing the impact of frugivores on germination is between gut-passed seeds and seeds remaining within intact fruit. We found that experiments involving comparisons between gut-passed and manually de-pulped seeds had substantially smaller effect sizes than those involving comparisons to whole/intact fruit. Although our meta-analysis showed large and statistically significant total effect sizes across all vertebrate groups, other recent taxon-specific meta-analyses have shown no statistically significant effect for bats (Saldaña-Vázquez *et al.* 2019) or small positive effects for neotropical primates (Fuzessy *et al.* 2016). However, these meta-analyses focused on studies that made comparisons either exclusively to manually de-pulped seeds or primarily to manually de-pulped seeds without accounting for study methods (Saldaña-Vázquez *et al.* 2019)(Fuzessy *et al.* 2016). By separately testing the effect of de-inhibition and scarification, we demonstrate that the total influence of bats and primates on germination is likely positive. These findings amplify previous calls encouraging researchers to include comparisons to seeds within whole fruit in order to characterize the total performance impacts of gut passage and to estimate the impacts of frugivore loss on plant populations (Samuels & Levey 2005; Costa-Pereira 2017; Fricke *et al.* 2019).

The experimental design of gut passage effect studies may offer an incomplete understanding of the role of fruit-frugivore interactions on germination for multiple reasons, in addition to the failure to test de-inhibition effects discussed above. First, most gut passage studies are conducted on fleshy-fruited plants with frugivores that are considered to be good seed dispersers. Seeds of species lacking fleshy fruit can germinate after ingestion by herbivores (Jaroszewicz *et al.* 2009; Lovas-Kiss *et al.* 2020). Such plant-animal interactions could be considered mutualistic with foliage serving as the reward to the animal partner (Janzen 1984). Likewise, animal species that are commonly understood to be herbivores or granivores often pass seeds of fleshy- and non-fleshy-fruited species intact and contribute to seed dispersal effectiveness (van Leeuwen *et al.* 2020). However, their total role in seed dispersal is poorly known because they are seldom the focus of study. Second, researchers typically report the proportion of gut-passed seeds that germinate, but use the number of intact seeds recovered from feces, rather than the number of seeds ingested, as the denominator. This causes a positive bias on estimated effect sizes when seeds are destroyed during gut passage. We recommend that future studies quantify the number of seeds ingested relative to number of seeds that pass intact and germinate to thoroughly characterize the effect of gut passage on germination. Future research that characterizes the total impacts from ingestion to germination can help generalize knowledge of plant species’ dependence on animals, and the importance of diverse animal vectors, for seed dispersal.

Our data synthesis spurs recommendations for taxa that should be prioritized for future study. We found that—despite the decades of relatively intensive experimental research to quantify gut passage effects—only a small portion of frugivorous birds and mammals have been tested. The ecological impacts of many vulnerable and endangered frugivore species are poorly known, especially among birds. The same knowledge gap exists for plants: Aslan et al (2013) estimated that 156,900 angiosperm species are vertebrate dispersed yet only 446 plant genera have been tested to determine how frugivore gut passage affects germination. These insights suggest that plant and animal species of conservation concern should be prioritized for future research. On the other hand, we note that measured effect sizes for a given frugivore species can be highly variable across experiments, even when the same plant-animal combination is tested. This suggests that individual studies on plant-animal pairs may only provide an approximate understanding of the functional importance of a given frugivore species. Thus, this meta-analysis and a future examination of the relationship between plant traits and the effects of gut passage may provide sufficient information to predict effects without conducting extensive labor-intensive experiments.

Trait-based approaches have potential for predicting gut passage effects on unstudied plant-animal combinations, and developing a quantitative understanding of gut passage effects across frugivores globally. Traits related to animal diet, body mass, and morphology of mouth and gut could predict gut passage effects by different frugivores. Fuzessy et al. (2016) showed substantial variation in effect sizes among neotropical primates explained by their primary diet and gut complexity. Functional traits of plants such as seed size, flesh-to-seed ratio, and shade tolerance may predict the benefits that plants receive from animal gut passage, and plant functional groups may be useful for predicting gut passage effects (Aslan *et al.* 2019; Rogers *et al.* 2021). A trait-based analysis of fruit consumed by fish found that fish are more likely to disperse fleshy-fruited species than dry-fruited species, but did not find a relationship between fruit traits related to color, shape, or size and the probability of dispersal (Correa *et al.* 2015). However, gut passage by fish is severely understudied, and the methods have been inconsistent (Costa-Pereira 2017), therefore limited conclusions can be drawn without additional data.

We found that plant or animal species included in the Global Invasive Species Database, indicating they are considered invasive in at least a portion of their range, do not differ in their gut passage effects from plants or animals that lack invasive populations. A priori, one could imagine multiple possible relationships between invasiveness and gut passage effects for plants. On one hand, plants with invasive populations may offer more flesh rewards to encourage seed dispersal, establishment, and expansion, and also be more dependent upon these dispersers for germination (Richardson et al. 2000). On the other hand, reduced dependence on mutualistic interactions may contribute to a plant’s ability to invade. Our analysis did not support either alternative, suggesting dependence on gut passage for germination is uncoupled from their propensity to be invasive. Invasive frugivore species exhibited similar impacts of gut passage on germination as non-invasive frugivores, suggesting that invasive frugivores also provide a similar quality of dispersal on average as their native counterparts (Vizentin-Bugoni *et al.* 2019). Overall, this suggests that traits may be more important than species origin in predicting gut passage effects.

The effects of frugivores on gut passage were more positive in temperate and subtropical locations than in the tropics, and on mainland systems than on islands. We caution that these effect sizes could be confounded by the study species targeted by researchers in these areas, yet find the patterns intriguing nonetheless. The increased benefit of gut passage in temperate and subtropical areas is surprising given that vertebrate seed dispersal is more common in tropical areas (Rogers *et al.* 2021). It is possible that the smaller number of temperate studies are more biased towards species most likely to benefit from dispersal. Alternatively, there may be limited successful life history strategies for fleshy-fruited species in temperate and subtropical areas compared to tropical areas. For example, tropical species may include many large-seeded plants with reduced dependence on frugivores for gut passage. Species on oceanic islands may be expected to exhibit smaller effect sizes because species with fewer dependencies on species interactions for reproduction and survival may be more likely to establish or persist in species-poor systems.

Because our meta-analysis shows that the total benefit of gut passage was large, the loss of these benefits may pose substantial demographic constraints for plant reproduction in ecosystems facing frugivore declines. For the average fleshy-fruited plant species studied, the mean effect size corresponds to more than a 60% reduction in germination probability for seeds not ingested by frugivores. We suggest that the loss of benefits of gut passage are underappreciated relative to other mechanisms that could negatively affect plant populations experiencing disperser loss. The loss of benefits associated with escape from conspecific negative distance- or density-dependent mortality (CNDD) is often highlighted as the primary negative consequence of seed disperser loss on plant populations. Yet a meta-analysis of experiments measuring the strength of CNDD (Comita et al. 2010) showed mean effect sizes, which were unrelated to study duration, corresponding to a roughly 25% reduction in survival for undispersed individuals. Although the effects of CNDD accrue over life stages, CNDD primarily impacts plant survival at the earliest life stages (Green & Harms 2017). The loss of gut passage benefits appears to be sizable relative to the loss of benefits associated with escape from CNDD.

Our meta-analysis brings renewed attention to a widespread yet under-appreciated ecological interaction. The simple act of removing flesh from a seed likely provides significant benefits for over half of the world’s plants (Aslan *et al.* 2013; Rogers *et al.* 2021). Many populations of avian, mammalian, reptilian, and fish seed dispersers are in decline in systems around the world; fewer individual frugivores, even in common species, means more fruits are left unconsumed and thus have a reduced chance of germination. When these condition-related benefits are combined with movement-related benefits of dispersal, the impacts on plant populations and communities are likely to be significant (Rogers *et al.* 2021). However, our finding that the primary benefits of frugivory come from de-inhibition rather than scarification, provides some optimism. First, most species do not require special treatment in the gut by a particular frugivore to germinate. Rather, any frugivore that consumes a given species and passes the seeds intact may confer some level of benefit, which increases the potential for compensation by remaining frugivores, even non-native species. Second, in the short-term and on an extremely limited spatial scale, humans may be able to maintain some plant species of conservation concern through fruit collection, manual de-pulping, and seed sowing. However, restoring fruit-frugivore mutualisms through rewilding will be necessary to restore this ecological function at larger taxonomic and geographic scales.

