## Supplemental Materials for "Frugivore gut passage increases seed germination: an updated meta-analysis"

### SUPPLEMENTARY MATERIALS

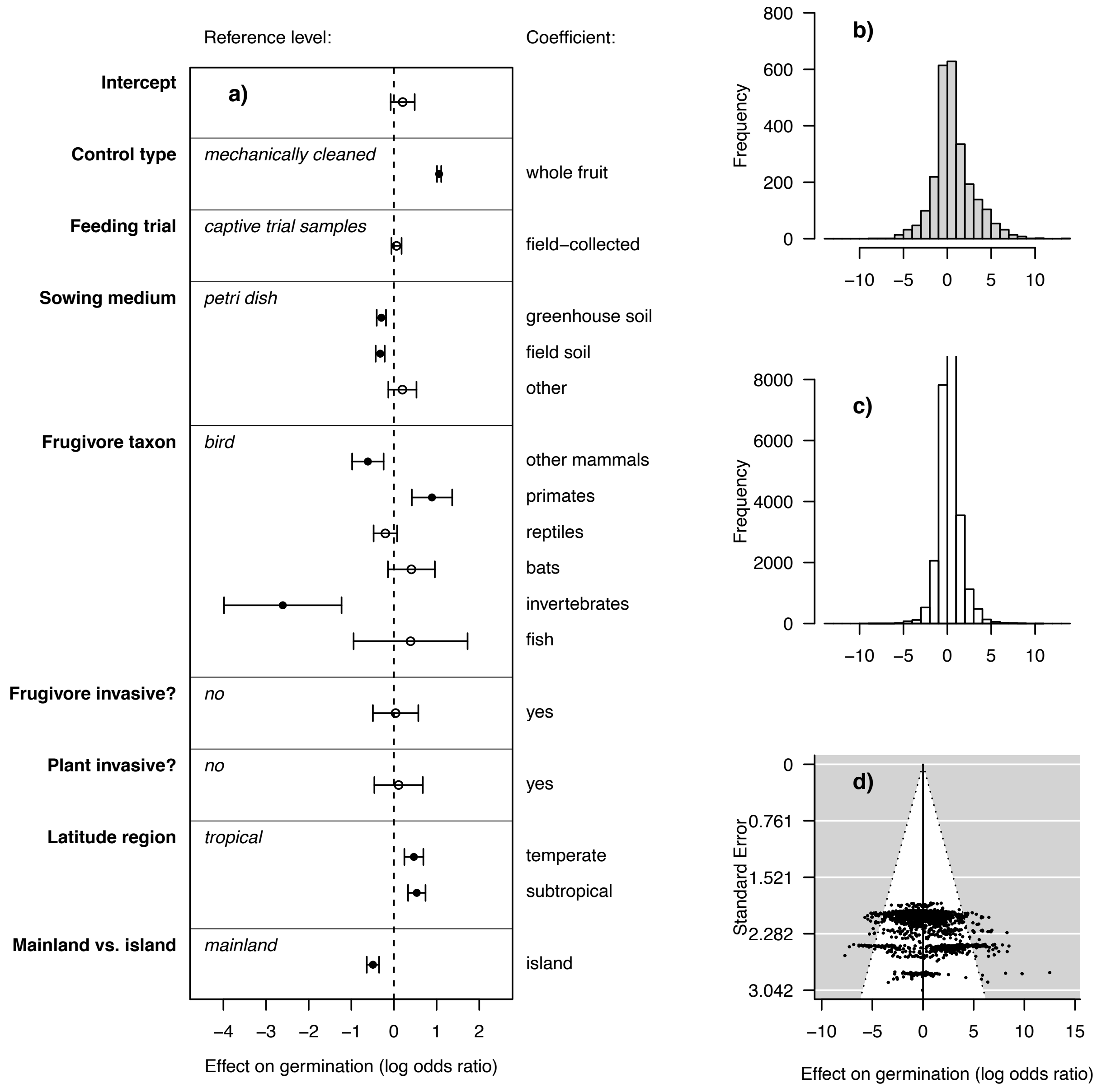


**Figure S1.** Summary of meta-analysis of experiments testing the impact of gut passage on seed germination. a) Log odds ratio effect sizes are shown for a full model including all predictor variables. The reference level for the categorical predictor variables are shown in italics, and effect sizes represent the effect associated with each coefficient described at the right. b) Frequency histogram of the unweighted effect of germination for all experiments included in the analysis. c) Histograms of effects weighted by in the inverse of variance. d) Funnel plot with each point showing the log odds ratio and standard error for each experiment.

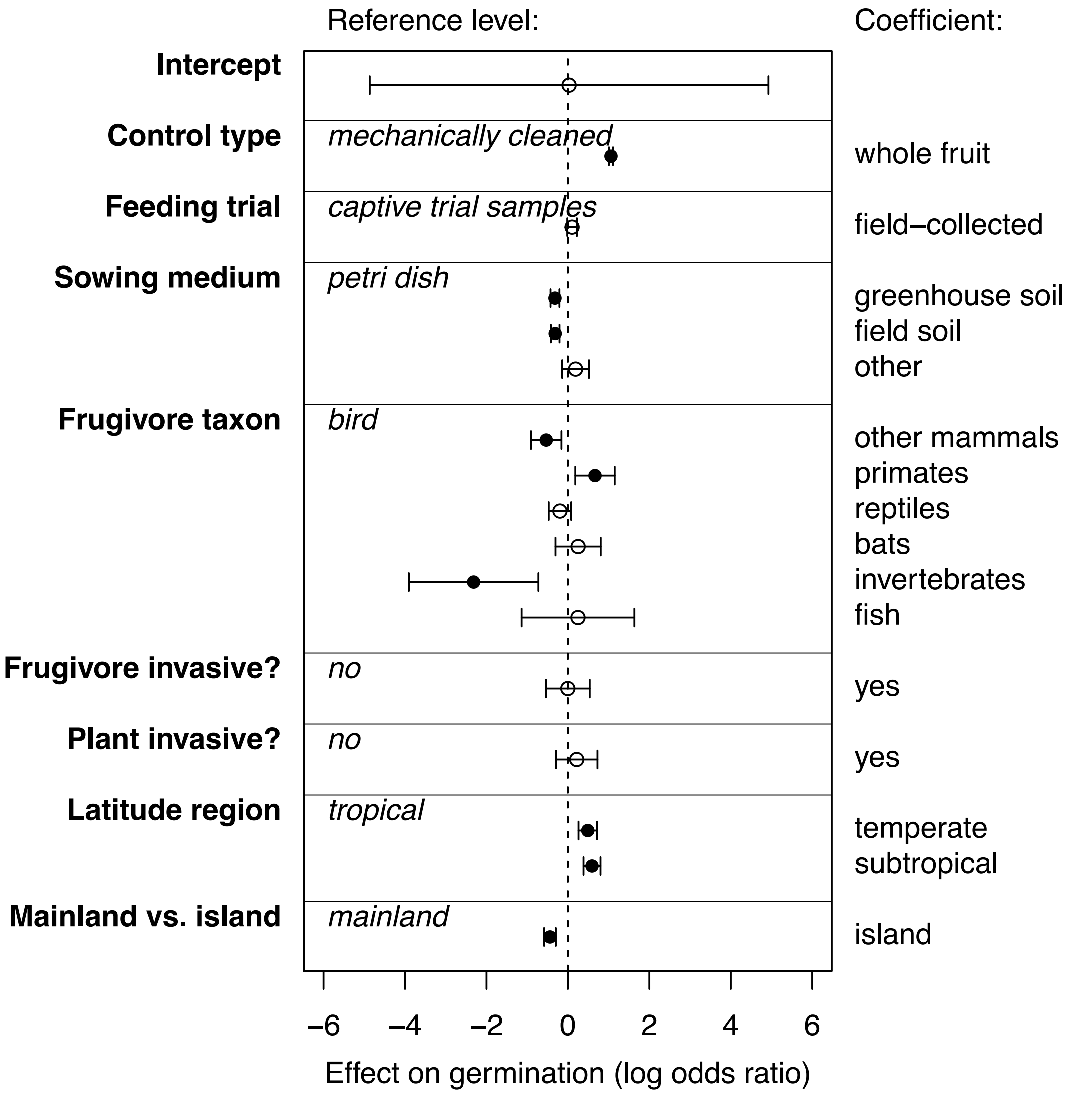


**Figure S2.** Summary of meta-analysis of experiments testing the impact of gut passage on germination incorporating plant phylogenetic non-independence.

**Table S1:** List of references used in the meta-analysis.

| Ref # | Reference |
|---|---|
| 1 | Agame, M. (1986) The role of mallard (Anas platyrhynchos) in distribution and germination of Najas marina L. seeds.. Oecologia. |
| 2 | Agami, M. (1988) The role of fish in distribution and germination of seeds of the submerged macrophytes Najas marina L. and Ruppia maritima L.. Oecologia. |
| 3 | Agmen, F. (2010) Seed dispersal by tantalus monkeys (Chlorocebus tantalus tantalus) in a Nigerian montane forest. African Journal of Ecology. |
| 4 | Alber, A. (2013) Frugivory and Seed Dispersal by Northern Pigtailed Macaques (Macaca leonina), in Thailand. International Journal of Primatology. 170-193. |
| 5 | Alves-Costa, C. (2007) Seed dispersal services by coatis (Nasua nasua, Procyonidae) and their redundancy with other frugivores in southeastern Brazil. 77-92. |
| 6 | Anderson, J. (2009) High-Quality Seed Dispersal by Fruit-Eating Fishes in Amazonian Floodplain Habitats. Oecologia. |
| 7 | Andriantsaralaza, S. (2014) The role of extinct giant tortoises in the germination of extant baobabAdansonia rubrostipa seeds in Madagascar. African Journal of Ecology. |
| 8 | Auger, J. (2002) Are American Black Bears (Ursus Americanus) Legitimate Seed Dispersers for Fleshy-Fruited shrubs?. The American Midland Naturalist. |
| 9 | Barnea, A. (1990) Differential germination of two closely related species of Solanum in response to bird ingestion.. Oikos. |
| 10 | Barnea, A. (1991) Does ingestion by birds affect seed germination?. Functional Ecology. 394-402. |
| 11 | Barnea, A. (1992) Effect of frugivorous birds on seed dispersal and germination of multi-seeded fruits. Acta Oecologica. 209-219. |
| 12 | Bas, J. (2006) Exclusive frugivory and seed dispersal of Rhamnus alaternus in the bird breeding season. Plant Ecology. |
| 13 | Beaune (2013) The Bonobo√¢‚Ç¨‚ÄúDialium Positive Interactions: Seed Dispersal Mutualism. American Journal of Primatology. |
| 14 | Ben√É¬≠tez-Malvido, J. (2003) Germination and Seed Damage in Tropical Dry Forest Plants Ingested by Iguanas. Journal of Herpetology. |
| 15 | Blake S (2012) Seed dispersal by Galapagos tortoises. Journal of Biogeography. |
| 16 | Blattmann, T. (2013) Gastropod Seed Dispersal: An Invasive Slug Destroys Far More Seeds in Its Gut than Native Gastropods. PLOS one. 8. |
| 17 | Boone, M. (2015) A Test of Potential Pleistocene Mammal Seed Dispersal in Anachronistic Fruits using Extant Ecological and Physiological Analogs. Southeastern Naturalist. |
| 18 | Borchert, M. (2011) Desiccation Sensitivity and Heat Tolerance of Prunus ilicifolia Seeds Dispersed by American Black Bears (Ursus americanus). Western North American Naturalist. |
| 19 | Bradford, M. (2010) Consequences of southern cassowary (Casuarius casuarius, L.) gut passage and deposition pattern on the germination of rainforest seeds. Austral Ecology. 325-333. |
| 20 | Braun, J. (1987) Box turtles (Terrapene carolina) as potential agents for seed dispesal. The American Midland Naturalist. 117. 312-318. |
| 21 | Bravo (2009) Implications of Behavior and Gut Passage for Seed Dispersal Quality: The Case of Blackand Gold Howler Monkeys. Biotropica. |
| 22 | Bravo, C. (2014) Effects of great bustard (Otis tarda) gut passage on black nightshade (Solanumnigrum) seed germination. Seed Science Research. |
| 23 | Brunner, H. (1976) A note on the dispersal of seeds of blackberry (Rubus procerus P. J. Muell.) by foxes and emus. Weed Research. |
| 24 | Bustamante, R. (1992) Are foxes legitimate and efficient seed dispersers? A field test. Acta Oecologia. 13. 203-208. |

| | |
|---|---|
| 25 | Calvino-Cancela, M. (2004) Ingestion and dispersal: direct and indirect effects of frugivores on seed viability and germination of Corema album(Empetraceae). Acta Oecologica. |
| 26 | Campos-Arceiz (2012) Asian tapirs are no elephans when it comes to seed dispersal. Biotropica. |
| 27 | Campos-Arceiz, Ahimsa (2008) Behavior rather than diet mediates seasonal differences in seed dispersal by Asian elephants. Ecology. 2684-2691. |
| 28 | Cantor, M. (2010) Potential seed dispersal by Didelphis albiventris(Marsupialia, Didelphidae) in highly disturbed environment. Biota Neotrop. |
| 29 | Carvalho-Ricardo, M. (2014) Frugivory and the effects of ingestion by bats on the seed germination of three pioneering plants. Acta Oecologica. 51-57. |
| 30 | Castilla, A. (2000) Does passage time through the lizard Podarcis lilfordiv¢,Ç¨,Ñ¢s guts affect germination performance in the plant Withania frutescens?. Acta Oecologica. |
| 31 | Castilla, M. (2009) The lizard Podarcis lilfordi as a potential disperser of the Solanaceae plant Lycopersicon esculentum: Can legitimate dispersers indirectly promote plant invasions?. Munibe Ciencias Naturales. |
| 32 | Catenacci, L. (2009) Seed Dispersal by Golden-headed Lion Tamarins Leontopithecus chrysomelas in Southern Bahian Atlantic Forest, Brazil. Biotropica. 744-750. |
| 33 | Celedon-Neghme, C. (2008) Legitimate seed dispersal by lizards in an alpine habitat: The case of Berberis empetrifolia (Berberidaceae) dispersed by Liolaemus belii (Tropiduridae). Acta Oecologica. 33. 265-271. |
| 34 | Chang, S. (2012) Frugivory by Taiwan Barbets (Megalaima nuchalis) and the effects of deinhibition and scarification on seed germination. Canadian Journal of Zoology. 640-650. |
| 35 | Clout, M. (1991) Germination of miro (Prumnopitys ferruginea) seeds after consumption by New Zealand pigeons (Hemiphaga novaeseelandiae). New Zealand Journal of Botany. 25-28. |
| 36 | Cochrane (2003) The need to be eaten: Balanites wilsoniana with and without elephant seed-dispersal. Journal of Tropical Ecology. 579-589. |
| 37 | Cochrane, J (2005) Endozoochory and the Australian bluebell: consumption of Billardiera fusiformis (Labill.) Payer (Pittosporaceae) seeds by three mammal species at Two Peoples Bay Nature Reserve, Western Australia.. Journal of the Royal Society of Western Australia. |
| 38 | Colon, C. (2013) THE IMPACT OF GUT PASSAGE BY BINTURONGS (ARCTICTIS BINTURONG) ON SEED GERMINATION. THE RAFFLES BULLETIN OF ZOOLOGY. |
| 39 | Crossland, D. (1996) Berry consumption by the American Robin, Turdus migra-torius, and the subsequent effect on seed germination, plant vigour, and dispersal of the lowbush blueberry, Vaccinium angustifolium. The Canadian Field Naturalist. |
| 40 | D'Avila, G. (2010) The role of avian frugivores on germination and potential seed dispersal of the Brazilian Pepper Schinus terebinthifolius. Biota Neotrop.. |
| 41 | Dunstan (2013) Dietary characteristics of Emus (Dromaius novaehollandiae) in semi-arid New South Wales, Australia, and dispersal and germination of ingested seeds. EMU. 168-176. |
| 42 | Ellison, A. (1993) Seed and seedling ecology of neotropical Melastomataceae.. Ecology. |
| 43 | Engel, T. (1997) Seed dispersal and plant regeneration by snakes?. Ecotropica. 3. 33-41. |
| 44 | Fedriani, J. (2009) Functional diversity in fruit-frugivore interactions: a field experiment with Mediterranean mammals. Ecography. |
| 45 | Fedriani, J. (2011) Dangerous liaisons disperse the Mediterranean dwarf palm: fleshy-pulp defensive role against seed predators. Ecology. |
| 46 | Figueiredo (1995) Germination ecology of Ficus luschnathiana drupelets after bird and bat ingestion. Acta Oecologica. 71-75. |
| 47 | Figueiredo, R. (1993) Ingestion of Ficus enormis seeds by howler monkeys (Alouatta fusca) in Brazil: effects on seed germination. Journal of Tropical Ecology. |
| 48 | Figueiredo, R. (1997) Ecological aspects of the dispersal of a melastomataceae by marmosets and howler monkeys (Primates: Platyrrhini) in a semideciduous forest of Southeastern Brazil. Rev. Ecol.. 52. 3-8. |
| 49 | Figuerira, J. (1994) Saurochory in Melocactus violaceus (Cactacea). Biotropica. |
| 50 | Figueroa, J. (2002) Effects of bird ingestion on seed germination of four woody species of the temperate rainforest of ChilovÉ¬© island, Chile. Plant Ecology. |

| | |
|---|---|
| 51 | FUKUI, A. (1995) The Role of the Brown-Eared Bulbul Hypsypetes amaurotis As a Seed Dispersal Agent. Population Ecology. |
| 52 | Gervais, J. (1998) The Potential for Seed Dispersal by the Banana Slug (Ariolimax columbianus). The American Midland Naturalist. |
| 53 | Gimeno, I. (2002) Recruitment of two Opuntia species invading abandoned olive groves. Acta Oecologica. |
| 54 | Gonz√É¬°lez-Di Pierro (2011) Effects of the Physical Environment and Primate Gut Passage on the Early Establishment of Ampelocera hottlei Standley in Rain Forest Fragments. Biotropica. |
| 55 | Griffiths, C. (2011) Resurrecting Extinct Interactions with Extant Substitutes. Current Biology. |
| 56 | Guerrero, S. (1997) Influencia de uma ave neotropical (Turdus rufiventris Vieillot) sobre a germina√É¬ß√É¬£o das sementes da figueiraasi√É¬°tica (Ficus microcarpa Linn.f.). Biotemas. 10. 27-34. |
| 57 | Guerta, R.S. (2011) Bird frugivory and seed germination of Myrsine umbellata and Myrsine lancifolia (Myrsinaceae) seeds in a cerrado fragment in southeastern Brazil. Biota Neotrop.. |
| 58 | Heer, K. (2010) Effects of ingestion by neotropical bats on germination parameters of native free-standing and strangler figs (Ficus sp., Moraceae). Oecologia. 425-435. |
| 59 | Hickey, J. (1999) An Evaluation of a Mammalian Predator, Martes americana, as a Disperser of Seeds. Oikos. |
| 60 | Holthuijzen, A. (1985) The avian seed dispersal system of eastern red cedar (Juniperus virginiana). Canadian Journal of Botany. 63. 1508-1515. |
| 61 | Honkavaara, J. (2007) Avian seed ingestion changes germination patterns of bilberry, Vaccinium myrtillus. Annales Botanici Fennici. 8-17. |
| 62 | Horn, M. (1997) Evidence for dispersal of fig seeds by the fruit-eating characid fish Brycon guatemalensis Regan in a Costa Rican tropical rain forest. Oecologia. 259-264. |
| 63 | Hunter, J. (1989) Seed dispersal and germination of Enterolobium cyclocarpum (Jacq.) Griseb. (Leguminosae: Mimosoideae): are megafauna necessary?. Journal of Biogeography. 369-378. |
| 64 | Izhaki, I. (1990) The effect of some mediterranean scrubland frugivores upon germination patterns. Journal of Ecology. |
| 65 | Izhaki, I. (1995) The effect of bat (Rousettus aegyptiacus) dispersal on seed germination in eastern Mediterranean habitats. Oecologia. 335-342. |
| 66 | Jacomassa, F. (2010) Birds and bats diverge in the qualitative and quantitative components of seed dispersal of a pioneer tree. Acta Oecologica. 493-496. |
| 67 | Jerozolimski, A. (2009) Are tortoises important seed dispersers in Amazonian forests?. Oecologia. 517-528. |
| 68 | Jordaan, L. (2012) Comparison of germination rates and fruit traits of indigenousSolanum giganteum and invasive Solanum mauritianum in South Africa. South African Journal of Botany. |
| 69 | Jordaan, L. (2012) Wahlberg√¢‚Ç¨‚Ñ¢s epauletted fruit bat (Epomophorus wahlbergi) as a potential dispersal agent for fleshy-fruited invasive alien plants: effects of handling behaviour on seed germination. Biological invasions. |
| 70 | Julliot, C. (1995) Seed dispersal by red howling monkeys in the tropical rain forest of French Guiana. International Journal of Primatology. |
| 71 | Kaufmann, S. (1991) Adaptations for a Two-Phase Seed Dispersal System Involving Vertebrates and Ants in a Hemiepiphytic Fig (Ficus microcarpa: Moraceae). American Journal of Botany. |
| 72 | Koike, S. (2008) Fruit phenology of Prunus jamasakuraand the feeding habit of the Asiatic black bear as a seed disperser. Ecological Research. |
| 73 | Krefting, Laurits (1949) The Role of Some Birds and Mammals in Seed Germination. Ecological Monographs. |
| 74 | Kunz, B. (2008) The Role of the Olive Baboon (Papio anubis, Cercopithecidae) as Seed Disperser in a Savanna-Forest Mosaic of West Africa. Journal of Tropical Ecology. |
| 75 | Ladley, J. (1996) DISPERSAL, GERMINATION AND SURVIVAL OF NEW ZEALAND MISTLETOES (LORANTHACEAE): DEPENDENCE ON BIRDS. New Zealand Journal of Ecology. |
| 76 | LaFleur, N. (2009) Does frugivory by European starlings (Sturnus vulgaris) facilitate germination in invasive plants?1. The Journal of the Torrey Botanical Society. |
| 77 | Lapenta, MJ (2008) Frugivory and seed dispersal of golden lion tamarin (Leontopithecus rosalia (Linnaeus, 1766)) in a forest fragment in the Atlantic Forest, Brazil. Brazilian Journal of Biology. 241-249. |

| | |
|---|---|
| 78 | Lara, C. (2009) Provenance, guts, and fate: Field and experimental evidence in a host-mistletoe-bird system. Ecoscience. |
| 79 | Lehouck, V. (2011) Avian fruit ingestion differentially facilitates seed germination of four fleshy-fruited plant species of an Afrotropical forest. Plant Ecology and Evolution. 96-100. |
| 80 | Lessa, L (2013) Effects of gut passage on the germination of seeds ingested by didelphid marsupials in a neotropical savanna. Acta Botanica Brasicila. 519-525. |
| 81 | Levey, D. (1986) Methods of seed processing by birds and seed deposition patterns. Frugivores and Seed Dispersal. |
| 82 | Lewis, D. (1987) Fruiting patterns, seed germination, and distribution of Sclerocarya caffra in an elephant-inhabited woodland. Biotropica. |
| 83 | Lieberman, D. (1979) Seed Dispersal by Baboons in the Shai Hills, Ghana. Ecology. |
| 84 | Lieberman, D. (1987) Notes on Seeds in Elephant Dung from Bia National Park, Ghana. Biotropica. |
| 85 | Lieberman, M. (1986) An Experimental Study of Seed Ingestion and Germination in a Plant-Animal Assemblage in Ghana. Journal of Tropical Ecology. |
| 86 | Linnebjerg, J. (2009) Gut passage effect of the introduced red-whiskered bulbul (Pycnonotus jocosus) on germination of invasive plant species in Mauritius. Austral Ecology. 272-277. |
| 87 | Liu, H. (2004) Seed dispersal by the Florida box turtle (Terrapene carolina bauri) in pine rockland forests of the lower Florida Keys, United States. Oecologia. 539-546. |
| 88 | Livingston, R. (1972) Influence of birds, stones and soil on the establishment of pasture junipers, Juniperus communis, and red cedar, J. virginiana in New England pastures.. Ecology. |
| 89 | Logan, D. (2006) Germination of kiwifruit, Actinidia chinensis, after passage through Silvereyes, Zosterops lateralis. New Zealand Journal of Ecology. 407-411. |
| 90 | Lombardi, J. (1993) Seed dispersal of Solanum lycocarpum St. Hil. (Solanaceae) by the maned wolf, Chrysocyon brachyurus Illiger (Mammalia, Canidae).. Ciencia e Cultura. |
| 91 | Lorinda, A. (2011) The role of avian frugivores in germination of seeds of fleshy-fruited invasive alien plants. Biological invasions. 1917-1930. |
| 92 | Mancilla-Leytv́É¬≥n, J. (2013) Effects of rabbit gut passage on seed retrieval and germination of three shrub species. Basic and Applied Ecology. 585-592. |
| 93 | Mandujanoa, M. (1997) Dormancy and endozoochorous dispersal ofOpuntia rastreraseeds in the southern Chihuahuan Desert √¢Àú‚Ä†. Jouranl of Arid Environments. |
| 94 | MAS, A. (2008) Two thrush species as dispersers of Miconia prasina (Sw.) DC. (Melastomataceae): an experimental approach. Brazilian Journal of Biology. |
| 95 | McDiarmid, R. (1977) Dispersal of Stemmadenia donnell-smithii (Apocynaceae) by Birds. Biotropica. |
| 96 | Meyer, G. (1998) Influence of Seed Processing by Frugivorous Birds on Germination Success of Three North American Shrubs. The American Midland Naturalist. 140. 129-139. |
| 97 | Midya, S. (1991) The effect of birds upon germination of banyan (Ficus bengalensis) seeds. Journal of Tropical Ecology. 7. 537-538. |
| 98 | Moll, D. (1995) Evidence for a role in seed dispersal by two tropical herbivorous turtles. Biotropica. 27. 121-127. |
| 99 | Moolna, A. (2008) Preliminary observations indicate that giant tortoise ingestion improves seed germination for an endemic ebony species in Mauritius. African Journal of Ecology. 217-219. |
| 100 | Motta, J. (2002) The Frugivorous Diet of the Maned Wolf, Chrysocyon brachyurus, in Brazil: Ecology and Conservation. |
| 101 | NA (1978) Adaptations for animal dispersal of one-seed juniper seeds. Oecologia. 32. 333-339. |
| 102 | NA (1978) The biology and autecology of Nitraria L. in Australia. II. Seed germination, seedling establishment and response to salinity. Australian Journal of Ecology. 3. |
| 103 | NA (1983) Dispersal of fruit seeds by black bears. |
| 104 | NA (1984) Fruit eating and seed dispersal by howling monkeys (Alouatta palliata) in the tropical rain forest of Los Tuxtlas, Mexico. American Journal of Primatology. 6. |
| 105 | NA (1985) Role of alien and native birds in the dissemination of firetree ( Myrica faya Ait. - Myricaceae) and associated plants in Hawaii. Pacific Science. 39. 372-378. |
| 106 | NA (1986) Seed dispersal by pygmy chimpanzees (Pan paniscus): A preliminary report. Primates. 27. 441-447. |

| | |
|---|---|
| 107 | NA (1986) Some aspects of dry land afforestation in the Sudan with special reference to Acacia tortilis (Forsk.) Hayne, A. senegal Willd. and Prosopis chilensis (Molina) stuntz. Forest Ecology and Management. 16. |
| 108 | NA (1990) Seed production and dispersal in the non-native, invasive succulent Carpobrotus edulis (Aizoaceae) in coastal strand communities of central California. Journal of Applied Ecology. 27. 693-702. |
| 109 | NA (1992) Balanites wilsoniana: Elephant dependent dispersal?. Journal of Tropical Ecology. 8. 275-283. |
| 110 | NA (1993) Consumo de g√°lbulos de sabina (juniperus phoenicea ssp turbinata guss, 1891) y dispersi√≥n de semillas por el conejo (oryctolagus cuniculus L.) en el parque nacional de Do√±ana. Do√±ana Acta Vertebrata. |
| 111 | NA (1995) Um teste de germinacao em sementes dispersas por macacos-Aranha em Maraca, Roraima; Brasil. Studies on Neotropical Fauna and Environment. 30. 31-36. |
| 112 | NA (1996) Seed dispersal in an African fig tree: Birds as high quantity, low quality dispersers?. Journal of Biogeography. 23. 553-563. |
| 113 | NA (1997) Carnivorous mammals as seed dispersers of myrtus communis (Myrtaceae) in the mediterranean shrublands. Plant Biosystems. 131. 189-195. |
| 114 | NA (1997) Dispersal and germination of Prosopis flexuosa (Fabaceae) seeds by desert mammals in Argentina. Journal of Arid Environments. 35. 707-714. |
| 115 | NA (1997) Role of birds on the regenaration of the woody Boscia senegalensis (Pers.) Lam. in Sahelian savanna in North Senegal [Role des oiseaux sur la regenaration du ligneux Bscia senegalensis (Pers.) Lam. en savane Sahelienne au Nord Senegal]. Revue d'Ecologie (La Terre et la Vie). 52. 239-260. |
| 116 | NA (1997) The diet of Pteropus voeltzkowi, an endangered fruit bat endemic to Pemba Island, Tanzania. African Journal of Ecology. 35. 351-360. |
| 117 | NA (1998) Frugivory and seed dispersal by four species of primates in Madagascar's eastern rain forest. Biotropica. 30. 425-437. |
| 118 | NA (1999) Germination rates of tree seeds ingested by coyotes and raccoons. American Midland Naturalist. 142. 71-76. |
| 119 | NA (1999) Old World fruit bats can be long-distance seed dispersers through extended retention of viable seeds in the gut. Proceedings of the Royal Society B: Biological Sciences. 266. |
| 120 | NA (1999) Potential dispersal of Cecropia hololeuca by the common Opossum (Didelphis Aurita) in Atlantic forest, southeastern Brazil. Revue d'Ecologie (La Terre et la Vie). 54. 327-332. |
| 121 | NA (1999) Seed dispersal by common ravens Corvus corax among island habitats (Canarian Archipelago). Ecoscience. 6. 56-61. |
| 122 | NA (1999) The dispersal of fruits and seeds of Poison-ivy, Toxicodendron radicans, by Ruffed Grouse, Bonasa umbellus, and squirrels, Tamiasciurus hudsonicus and Sciurus carolinensis. Canadian Field-Naturalist. 113. 616-620. |
| 123 | NA (2000) Bacca quo vadis: Regeneration niche differences among seven sympatric Vaccinium species on headlands of Newfoundland. Seed Science Research. 10. 89-97. |
| 124 | NA (2000) Bet-hedging and germination in the Australian arid zone shrub Acacia ligulata. Austral Ecology. 25. 368-374. |
| 125 | NA (2000) Germination of seeds from three species dispersed by black howler monkeys (Alouatta caraya). Folia Primatologica. 71. 342-345. |
| 126 | NA (2000) Inhibition by pulp juice and enhancement by ingestion on germination of bird-dispersed Prunus seeds. Journal of Forest Research. 5. 213-215. |
| 127 | NA (2000) Mistletoe seed dispersal by a marsupial. Nature. 408. 929-930. |
| 128 | NA (2001) Fruit fate, seed germination and growth of an invasive vine - An experimental test of 'sit and wait' strategy. Biological Invasions. 3. |
| 129 | NA (2001) Seed dispersal by a diurnal primate community in the Dja Reserve, Cameroon. Journal of Tropical Ecology. 17. 787-808. |
| 130 | NA (2001) Seed dispersal: Directed deterrence by capsaicin in chillies. Nature. 412. 403-404. |
| 131 | NA (2002) A pest and an invader: White-tailed deer (Odocoileus virginianus Zimm.) as a Seed dispersal agent for honeysuckle shrubs (Lonicera L.). Natural Areas Journal. 22. 230-234. |
| 132 | NA (2002) Are predatory birds effective secondary seed dispersers?. Biological Journal of the Linnean Society. 75. |

| | |
|---|---|
| 133 | NA (2002) Contribution by possums to seed rain and subsequent seed germination in successional vegetation, Canterbury, New Zealand. New Zealand Journal of Ecology. 26. 121-128. |
| 134 | NA (2002) Feeding ecology of Pteropus rufus (Pteropodidae) in the littoral forest of Sainte Luce, SE Madagascar. Acta Chiropterologica. 4. 33-47. |
| 135 | NA (2002) Germination in Prosopis ferox seeds: Effects of mechanical, chemical and biological scarificators. Journal of Arid Environments. 50. 185-189. |
| 136 | NA (2002) Interacting effects of forest fragmentation and howler monkey foraging on germination and disperal of fig seeds. ORYX. 36. |
| 137 | NA (2002) Seed dispersal by the sloth bear (Melursus ursinus) in South India. Biotropica. 34. 474-477. |
| 138 | NA (2002) The lizard Teius teyou (Squamata: Teiidae) as a legitimate seed disperser in the dry Chaco forest of Argentina. Studies on Neotropical Fauna and Environment. 37. |
| 139 | NA (2002) The role of seed dispersers in the population dynamics of the columnar cactus Neobuxbaumia tetetzo. Ecology. 83. 2617-2629. |
| 140 | NA (2003) Effect of ingestion by five avian dispersers on the retention time, retrieval and germination of Ruppia maritima seeds. Functional Ecology. 17. |
| 141 | NA (2003) Effects of Passage Through Tamarin Guts on the Germination Potential of Dispersed Seeds. International Journal of Primatology. 24. |
| 142 | NA (2003) Erratum: Effect of ingestion by bats and birds on seed germination of Stenocereus griseus and Subpilocereus repandus (Cactaceae) (Journal of Tropical Ecology (2003) 19:1 (19-25)). Journal of Tropical Ecology. 19. |
| 143 | NA (2003) Frugivory and seed dispersal of Miconia urophylla (Melastomataceae) by birds in a fragment of secondary Atlantic forest in Juiz de Fora, Minas Gerais State, Brazil [Frugivoria e dispers√£o de sementes de Miconia urophylla (Melastomataceae) por aves em um fragmento de Mata Atl√¢ntica secund√°ria em Juiz de Fora, Minas Gerais, Brasil]. Ararajuba. 11. 173-180. |
| 144 | NA (2003) Observations on food habits of Asiatic black bear in Kedarnath wildlife sanctuary, India: Preliminary evidence on their role in seed germination and dispersal. Ursus. 14. 99-103. |
| 145 | NA (2004) Observations on the role of frugivorous bats as seed disperses in Costa Rican secondary humid forests. Acta Chiropterologica. 6. 111-119. |
| 146 | NA (2004) Relationships between alien plants and an alien bird species on Reunion Island. Journal of Tropical Ecology. 20. |
| 147 | NA (2005) Disruption of a plant-lizard seed dispersal system and its ecological effects on a threatened endemic plant in the Balearic Islands. Conservation Biology. 19. 421-431. |
| 148 | NA (2005) Frugivory and seed dispersal by foxes in relation to mammalian prey abundance in a semiarid thornscrub. Austral Ecology. 30. |
| 149 | NA (2005) Fruit consumption and seed dispersal of Dimorphandra mollis Benth. (Leguminosae) by the lowland tapir in the cerrado of Central Brazil.. Brazilian journal of biology = Revista brasleira de biologia.. 65. 407-413. |
| 150 | NA (2005) Invasional meltdown potential: Facilitation between introduced plants and mammals on French Mediterranean islands. Ecoscience. 12. 248-256. |
| 151 | NA (2005) Post-dispersal seed removal and germination selected tree species dispersed by Cebus capucinus on Barro Colorado Island, Panama. Biotropica. 37. 73-80. |
| 152 | NA (2006) Comparative seed dispersal effectiveness of sympatric Alouatta guariba and Brachyteles arachnoidesin southeastern Brazil. Biotropica. 38. |
| 153 | NA (2006) Effects of ingestion by cattle and immersion in hot water and acid on the germinability of rain tree (Albizia saman) seeds. Tropical Grasslands. 40. 244-253. |
| 154 | NA (2006) Germination of Ficus insipida (Moraceae) seeds from toucan (Ramphastos sulfuratus) and spider monkey (Ateles geoffroyi) feces [Germinaci√≥n de semillas de Ficus insipida (Moraceae) defecadas por tucanes (Ramphastos sulfuratus) y monos ara√±a (Ateles geoffroyi)]. Revista de Biologia Tropical. 54. 387-394. |
| 155 | NA (2006) Rabbits (Oryctolagus cuniculus) as dispersers of Retama monosperma seeds in a coastal dune system. Ecoscience. 13. |
| 156 | NA (2006) Red fox (Vulpes vulpes L.) favour seed dispersal, germination and seedling survival of Mediterranean Hackberry (Celtis australis L.). |

| | |
|---|---|
| 157 | NA (2006) Reproductive ecology of invasive Ochna serrulata (Ochnaceae) in south-eastern Queensland. Australian Journal of Botany. 54. |
| 158 | NA (2006) Restoring dry Afromontane forest using bird and nurse plant effects: Direct sowing of Olea europaea ssp. cuspidata seeds. Forest Ecology and Management. 230. |
| 159 | NA (2006) Ruminant-mediated seed dispersal of an economically valuable tree in Indian dry forests. Biotropica. 38. 679-682. |
| 160 | NA (2006) The effect of seed morphology on the potential dispersal of aquatic macrophytes by the common carp (Cyprinus carpio). Freshwater Biology. 51. 2063-2071. |
| 161 | NA (2007) Decreased frugivory and seed germination rate do not reduce seedling recruitment rates of Aristotelia chilensis in a fragmented forest. Biodiversity and Conservation. 16. |
| 162 | NA (2007) European pond turtles (Emys orbicularis) as alternative dispersers of "water-dispersed" waterlily (Nymphaea alba). Ecoscience. 14. |
| 163 | NA (2007) Frugivory and seed dispersal by Asian elephants, Elephas maximus, in a moist evergreen forest of Thailand. Journal of Tropical Ecology. 23. |
| 164 | NA (2007) Germination in seed species ingested by opossums: Implications for seed dispersal and forest conservation. Brazilian Archives of Biology and Technology. 50. 921-928. |
| 165 | NA (2007) Germination rate and velocity of seeds of Cecropia pachystachya Tr√©cul (Cecropiaceae) eaten by Artibeus lituratus (Olfers, 1818) (Chiroptera: Phyllostomidae) [Taxa e velocidade de germina√ß√£o de sementes de Cecropia pachystachya Tr√©cul (Cecropiaceae) ingeridas por Artibeus lituratus (Olfers, 1818) (Chiroptera: Phyllostomidae)]. Acta Scientiarum - Biological Sciences. 29. 395-399. |
| 166 | NA (2007) Predicting spatial patterns of plant recruitment using animal-displacement kernels. PLoS ONE. 2. |
| 167 | NA (2007) Reproductive versus vegetative recruitment of the invasive tree Schinus terebenthifolius: Implications for restoration on Reunion Island. Restoration Ecology. 15. |
| 168 | NA (2007) Secondary seed dispersal systems, frugivorous lizards and predatory birds in insular volcanic badlands. Journal of Ecology. 95. 1394-1403. |
| 169 | NA (2007) The feeding ecology of Eidolon dupreanum (Pteropodidae) in eastern Madagascar. African Journal of Ecology. 45. |
| 170 | NA (2008) Ecological efficiency and legitimacy in seed dispersal of an endemic shrub (Lithrea caustica) by the European rabbit (Oryctolagus cuniculus) in central Chile. Journal of Arid Environments. 72. |
| 171 | NA (2008) Endozoochory by native and exotic herbivores in dry areas: Consequences for germination and survival of Prosopis seeds. Seed Science Research. 18. |
| 172 | NA (2008) Frugivorous bats (Mammalia, Chiroptera) in Cecropia pachystachya (Urticaceae) and their effects in seed germination [Frugivoria de morcegos (Mammalia, Chiroptera) em Cecropia pachystachya (Urticaceae) e seus efeitos na germina√ß√£o das sementes]. Papeis Avulsos de Zoologia. 48. |
| 173 | NA (2008) Frugivory and seed dispersal by the yellow-throated marten, Martes flavigula, in a subtropical forest of China. Journal of Tropical Ecology. 24. |
| 174 | NA (2008) Potential role of frugivorous birds (Passeriformes) on seed dispersal of six plant species in a restinga habitat, southeastern Brazil. Revista de Biologia Tropical. 56. 205-216. |
| 175 | NA (2008) Seed dispersal and establishment of endangered plants on oceanic islands: The Janzen-Connell model, and the use of ecological analogues. PLoS ONE. 3. |
| 176 | NA (2009) Germination patterns throughout an insular altitudinal gradient: The case of the Macaronesian endemic plant Rubia fruticosa Ait. (Rubiaceae) in El Hierro (Canary Islands). Flora: Morphology, Distribution, Functional Ecology of Plants. 204. |
| 177 | NA (2009) Importance of a tropical tree (Dendropanax arboreus) for Neartic migrant birds in Mexico [Importance de un arbol tropical (dendropanax arboreus) para aves migratorias ne√°rticas en m√©xico]. Ornitologia Neotropical. 20. 391-399. |
| 178 | NA (2009) Reproductive ecology of the endangered enigmatic mauritian endemic Roussea simplex (rousseaceae). International Journal of Plant Sciences. 170. 42-52. |
| 179 | NA (2009) Seed dispersal effectiveness by understory birds on Dendropanax arboreus in a fragmented landscape. Biodiversity and Conservation. 18. 3357-3365. |

| | |
|---|---|
| 180 | NA (2009) Seed dispersal of Solanum thomasiifolium Sendtner (Solanaceae) in the Linhares Forest, Esp√≠rito Santo state, Brazil [(Dispers√£o de sementes de Solanum thomasiifolium Sendtner (Solanaceae) na Floresta de Linhares, Esp√≠rito Santo, Brasil]. Acta Botanica Brasilica. 23. 1171-1179. |
| 181 | NA (2009) The Crab-eating Fox (Cerdocyon thous) as a secondary seed disperser of Eugenia umbelliflora (Myrtaceae) in a Restinga forest of southeastern Brazil [O chachorro do mato (Cerdocyon thous) como dispersor secund√°rio de Eugenia umbelliflora (Myrtaceae) em uma floresta de Restinga no sudeste do Brasil]. Biota Neotropica. 9. 271-274. |
| 182 | NA (2009) The key role of a ring ouzel tardus torquatus wintering population in seed dispersal of the endangered endemic juniperus cedrus in an insular environment. Acta Ornithologica. 44. |
| 183 | NA (2009) The potential key seed-dispersing role of the arboreal marsupial Dromiciops gliroides. Acta Oecologica. 35. 8-13. |
| 184 | NA (2010) Comparative seed and dispersal ecology of three exotic subtropical asparagus species. Invasive Plant Science and Management. 3. |
| 185 | NA (2010) Effect of gut passage and dung on seed germination and seedling growth: donkeys and a multipurpose mesquite from a Mexican inter-tropical desert. Journal of Arid Environments. 74. |
| 186 | NA (2010) Effect of red ruffed lemur gut passage on the germination of native rainforest plant species. Lemur News. 15. 39-42. |
| 187 | NA (2010) Postdispersal removal and germination of seed dispersed by cercopithecus nictitans in a west African montane forest. Folia Primatologica. 81. |
| 188 | NA (2010) Potencial dispers«éo de sementes por Didelphis albiventris (Marsupialia, Didelphidae) em ambiente altamente perturbado [Potential seed dispersal by Didelphis albiventris (Marsupialia, Didelphidae) in highly disturbed environment]. Biota Neotropica. 10. 45-51. |
| 189 | NA (2010) Seed dispersal by red-eared sliders (Trachemys scripta elegans) and common snapping turtles (Chelydra serpentina). Chelonian Conservation and Biology. 9. 289-294. |
| 190 | NA (2011) Avian consumption and seed germination of the hemiparasitic mistletoe Agelanthus natalitius (Loranthaceae). Journal of Ornithology. 152. 643-649. |
| 191 | NA (2011) Body size determines rates of seed dispersal by giant king crickets. Population Ecology. 53. |
| 192 | NA (2011) Differential seed dispersal systems of endemic junipers in two oceanic Macaronesian archipelagos: The influence of biogeographic and biological characteristics. Plant Ecology. 212. |
| 193 | NA (2011) Differential seed handling by two African primates affects seed fate and establishment of large-seeded trees. Acta Oecologica. 37. 578-586. |
| 194 | NA (2011) Effectiveness of Spider Monkeys (Ateles geoffroyi vellerosus) as Seed Dispersers in Continuous and Fragmented Rain Forests in Southern Mexico. International Journal of Primatology. 32. 177-192. |
| 195 | NA (2011) Frugivory and potential seed dispersal by the marsupial Gracilinanus agilis (Didelphidae: Didelphimorphia) in areas of Cerrado in central Brazil [Frugivoria e potencial dispers√£o de sementes pelo marsupial gracilinanus agilis (Didelphidae: Didelphimorphia) em √°reas de cerrado no brasil central]. Biotropica. 45. 465-473. |
| 196 | NA (2011) Ingestion by a freshwater turtle alters germination of bottomland hardwood seeds. Wetlands. 31. |
| 197 | NA (2011) Legitimate seed dispersal Ugni molinae Turcz. (Myrtaceae), by monito del monte, Dromiciops gliroides [Leg√≠tima dispersi√≥n de semillas Ugni molinae Turcz. (Myrtaceae), por monito del monte, Dromiciops gliroides]. Gayana - Botanica. 68. 309-312. |
| 198 | NA (2011) Plant-ungulate interaction: Goat gut passage effect on survival and germination of Mediterranean shrub seeds. Journal of Vegetation Science. 22. |
| 199 | NA (2011) Seed dispersal by megaherbivores: Do Asian elephants disperse Mallotus philippinensis, a main food tree in northern India and Nepal?. Journal of Natural History. 45. 915-921. |
| 200 | NA (2011) Seed dispersal by the Indian grey hornbill Ocyceros Birostris in Eastern Ghats, India. Ecotropica. 17. 71-77. |
| 201 | NA (2011) Successful germination of seeds following passage through orang-utan guts. Journal of Tropical Ecology. 27. |
| 202 | NA (2012) Black bears Ursus americanus are effective seed dispersers, with a little help from their friends. Oikos. 121. 589-596. |

| | |
|---|---|
| 203 | NA (2012) Can rodents enhance germination rates in rainforest seeds?. Ecological Management and Restoration. 13. |
| 204 | NA (2012) Diet and seed dispersal: Does the Colombian Chachalaca (Ortalis Columbiana) affect germination of Ingested seeds? [Dieta y dispersi√≥n de semillas: ¬øAfecta la Guacharaca Colombiana (Ortalis Columbiana) la germinaci√≥n de las semillas Consumidas?]. Ornitologia Neotropical. 23. 439-453. |
| 205 | NA (2012) Differential seed germination of a keystone palm (Euterpe edulis) dispersed by avian frugivores. Journal of Tropical Ecology. 28. 615-618. |
| 206 | NA (2012) Dispersion of Melocactus glaucescens and M. paucispinus (Cactaceae) in the municipality of Morro do Chap√©u, Chapada Diamantina - BA [Dispers√£o de sementes de Melocactus glaucescens e M. paucispinus (Cactaceae), no Munic√≠pio de Morro do Chap√©u, Chapada Diamantina - BA]. Acta Botanica Brasilica. 26. 481-492. |
| 207 | NA (2012) Effective dispersal of large seeds by Baird's tapir: A large-scale field experiment. Journal of Tropical Ecology. 28. 119-122. |
| 208 | NA (2012) Recovery and germination of dichrostachys cinerea seeds fed to goats (Capra hircus). Rangeland Ecology and Management. 65. |
| 209 | NA (2012) Seed dispersal by a captive corvid: The role of the 'Alala'(Corvus hawaiiensis) in shaping Hawai'i's plant communities. Ecological Applications. 22. 1718-1732. |
| 210 | NA (2012) Seed dispersal of matai (Prumnopitys taxifolia) by feral pigs (Sus scrofa). New Zealand Journal of Ecology. 36. |
| 211 | NA (2012) Significance and extent of secondary seed dispersal by predatory birds on oceanic islands: The case of the Canary archipelago. Journal of Ecology. 100. |
| 212 | NA (2012) Species-specific outcomes of avian gut passage on germination of Melastomataceae seeds. Plant Ecology and Evolution. 145. |
| 213 | NA (2013) Antelope mating strategies facilitate invasion of grasslands by a woody weed. Oikos. 122. |
| 214 | NA (2013) Contrasting Selective Pressures on Seed Traits of Two Congeneric Species by Their Main Native Guilds of Dispersers on Islands. PLoS ONE. 8. |
| 215 | NA (2013) Didelphis albiventris as germination inductors of Rapanea ferruginea (Myrcinaceae) in Cerrado area in Mato Grosso do Sul, Brazil [Didelphis albiventris como indutor de germina√ß√£o de Rapanea ferruginea (Myrcinaceae) em √°rea de cerrado, Mato Grosso do Sul, Brasil]. Iheringia - Serie Zoologia. 103. |
| 216 | NA (2013) Ecological restoration and reforestation of fragmented forests in kianjavato, Madagascar. International Journal of Ecology. |
| 217 | NA (2013) Ecological services performed by the bonobo (Pan paniscus): Seed dispersal effectiveness in tropical forest. Journal of Tropical Ecology. 29. 367-380. |
| 218 | NA (2013) Endozoochorus seed dispersal by goats: Recovery, germinability and emergence of five Mediterranean shrub species. Spanish Journal of Agricultural Research. 11. |
| 219 | NA (2013) Frugivory and seed dispersal by Tropidurus torquatus (Squamata: Tropiduridae) in southern Brazil. Herpetological Journal. 23. 75-79. |
| 220 | NA (2013) Specialized Seed Dispersal in Epiphytic Cacti and Convergence with Mistletoes. Biotropica. 45. 465-473. |
| 221 | NA (2013) The role of domestic goats in the conservation of four endangered species of cactus: Between dispersers and predators. Applied Vegetation Science. 16. |
| 222 | NA (2014) Effects of fruit ingestion by Taiwan Barbet (Megalaima nuchalis) on seed germination of four native tree species in Taiwan. Taiwan Journal of Forest Science. 29. |
| 223 | NA (2014) Russian olive (Elaeagnus angustifolia) dispersal by European starlings (Sturnus vulgaris). Invasive Plant Science and Management. 7. 425-431. |
| 224 | NA (2014) Seed dispersal by rhesus macaques Macaca mulatta in Northern India. American Journal of Primatology. 76. |
| 225 | NA (2014) Seed source, seed traits, and frugivore habits: Implications for dispersal quality of two sympatric primates. American Journal of Botany. 101. |
| 226 | NA (2014) The Role of the White-Winged Guan (Penelope albipennis) in Seed Dispersal and Predation in Tumbesian Dry Forest, Peru. Journal of Sustainable Forestry. 33. 184-194. |

| | |
|---|---|
| 227 | NA (2015) Avian‚ÄìShrub interactions: Ingestion, seed recovery and germination of three Mediterranean shrub species fed to quail (Coturnix coturnix). Russian Journal of Ecology. 46. |
| 228 | NA (2015) Dispersal of banana passionfruit (Passiflora tripartita var. mollissima) by exotic mammals in new zealand facilitates plant invasiveness. New Zealand Journal of Ecology. 39. 43-49. |
| 229 | NA (2015) Do animal‚Äìplant interactions influence the spatial distribution of Aristotelia chilensis shrubs in temperate forests of southern South America?. Plant Ecology. 216. 383-394. |
| 230 | NA (2015) Does the passage of seeds through frugivore gut affect their storage: A case study on the endangered plant Euryodendron excelsum. Scientific Reports. 5. |
| 231 | NA (2015) Frugivory by native (Lycalopex spp.) and allochthonous (Canis lupus familiaris) canids reduces the seed germination of litre (Lithrea caustica) in central Chile [La frugivor√≠a por c√°nidos nativos (Lycalopex spp.) y al√≥ctonos (Canis lupus familiaris) reduce la germinaci√≥n de semillas de litre (lithrea caustica) en Chile central]. Bosque. 36. |
| 232 | NA (2015) Fruit and seed traits of the elephant-dispersed African savanna plant Balanites maughamii. Journal of Tropical Ecology. 31. |
| 233 | NA (2015) In the elephant's seed shadow: The prospects of domestic bovids as replacement dispersers of three tropical Asian trees. Ecology. 96. 2093-2105. |
| 234 | NA (2015) Seed dispersal and predation of Buchenavia tomentosa Eichler (Combretaceae) in a Cerrado sensu stricto, midwest Brazil [Dispers√£o e preda√ß√£o de sementes de Buchenavia tomentosa Eichler (Combretaceae) em Cerrado sentido restrito, centro-oeste do Brasil]. Brazilian Journal of Biology. 75. |
| 235 | NA (2015) The potential for birds to disperse the seeds of Acacia cyclops, an invasive alien plant in South Africa. Ibis. 157. |
| 236 | NA (2015) The unnoticed effect of a top predator on complex mutualistic ecological interactions. Biological Invasions. 17. 1655-1665. |
| 237 | NA (2016) Are birds, wind and gravity legitimate dispersers of fleshy-fruited invasive plants on Robinson Crusoe Island, Chile?. Flora: Morphology, Distribution, Functional Ecology of Plants. 224. 167-171. |
| 238 | NA (2016) Asiatic Callosciurus squirrels as seed dispersers of exotic plants in the Pampas. Current Zoology. 62. |
| 239 | NA (2016) Bird and ant synergy increases the seed dispersal effectiveness of an ornithochoric shrub. Oecologia. 181. 507-518. |
| 240 | NA (2016) Diet and effects of Sanford‚Äôs brown lemur (Eulemur sanfordi, Archbold 1932) gut-passage on the germination of plant species in Amber forest, Madagascar. Zoological Studies. 55. |
| 241 | NA (2016) Diet of the endangered big-headed turtle Platysternon megacephalum. PeerJ. 2016. 10-NA. |
| 242 | NA (2016) Does seed coat structure modulate gut-passage effects on seed germination? Examples from Miconieae DC. (Melastomataceae). Seed Science Research. 26. |
| 243 | NA (2016) Ecosystem services provided by a large endangered primate in a forest-savanna mosaic landscape. Biological Conservation. 203. 55-66. |
| 244 | NA (2016) Effects of invasive Green Iguanas (Iguana iguana) on seed germination and seed dispersal potential in southeastern Puerto Rico. Biological Invasions. 18. |
| 245 | NA (2016) Introducing cultivated trees into the wild: Wood pigeons as dispersers of domestic olive seeds. Acta Oecologica. 71. |
| 246 | NA (2016) Myiarchus flycatchers are the primary seed dispersers of Bursera longipes in a Mexican dry forest. PeerJ. 2016. |
| 247 | NA (2016) Possible implications of seed dispersal by the howler monkey for the early recruitment of a legume tree in small rain-forest fragments in Mexico. Journal of Tropical Ecology. 32. |
| 248 | NA (2016) Seed dispersal effectiveness: A comparison of four bird species feeding on seeds of invasive Acacia cyclops in South Africa. South African Journal of Botany. 105. 259-263. |
| 249 | NA (2016) Seed dispersal potential of Asian elephants. Acta Oecologica. 77. |
| 250 | NA (2016) Seed removal by lizards and effect of gut passage on germination in a columnar cactus of the Caatinga, a tropical dry forest in Brazil. Journal of Arid Environments. 135. |
| 251 | NA (2016) Size-based fruit selection by a keystone avian frugivore and effects on seed viability. New Zealand Journal of Botany. 1-16. |

| | |
|---|---|
| 252 | NA (2016) The austral thrush (Turdus falcklandii) reduces the seed germination for the urban ornamental tree glossy privet (Ligustrum lucidum) in Santiago, Chile. Revista Brasileira de Botanica. 39. |
| 253 | NA (2017) Comparative effects of native frugivores and introduced rodents on seed germination in New-Caledonian rainforest plants. Biological Invasions. 19. 351-363. |
| 254 | NA (2017) Does avian gut passage favour seed germination of woody species of the Chaco serrano Woodland in Argentina?. Botany. 95. NA-493. |
| 255 | NA (2017) Frugivory and seed dispersal effectiveness in two Miconia (Melastomataceae) species from ferruginous campo rupestre. Seed Science Research. 27. NA-65. |
| 256 | NA (2017) Generalist dispersers promote germination of an alien fleshy-fruited tree invading natural grasslands. PLoS ONE. 12. |
| 257 | NA (2017) Germination and clonal propagation of the endemic shrub Corema album, a vulnerable species with conservation needs and commercial interest. Natural Product Communications. 12. NA-267. |
| 258 | NA (2017) Neglected seed dispersers: endozoochory by Javan lutungs (Trachypithecus auratus) in Indonesia. Biotropica. 49. NA-539. |
| 259 | NA (2017) Passage through artibeus lituratus (Olfers, 1818) increases germination of cecropia pachystachya (Urticaceae) seeds. Tropical Conservation Science. 10. |
| 260 | NA (2017) Preference of an insular flying fox for seed figs enhances seed dispersal of a dioecious species. Biotropica. 49. NA-511. |
| 261 | NA (2017) Quantifying seed dispersal by birds and possums in a lowland New Zealand forest. New Zealand Journal of Ecology. 41. |
| 262 | NA (2017) Seed dispersal effectiveness of the western lowland gorilla (Gorilla gorilla gorilla) in Gabon. African Journal of Ecology. |
| 263 | NA (2017) Size-based fruit selection by a keystone avian frugivore and effects on seed viability. New Zealand Journal of Botany. 55. NA-118. |
| 264 | NA (2017) Synchronous fruiting and common seed dispersers of two endemic columnar cacti in the Caatinga, a dry forest in Brazil. Plant Ecology. 218. NA-1325. |
| 265 | NA (2017) The importance of austral thrush Turdus falcklandii in seed germination of pitra Myrceugenia planipes [La importancia del zorzal austral Turdus falcklandii en la germinaci√≥n de semillas de pitra Myrceugenia planipes]. Revista Mexicana de Biodiversidad. 88. NA-474. |
| 266 | Nakamoto, A. (2008) Feeding effects of Orii's flying-fox (Pteropus dasymallus inopinatus) on seed germination of subtropical trees on Okinawa-jima Island. Tropics. |
| 267 | Nchanji, A. (2003) Seed germination and early seedling establishment of some elephant-dispersed species in Banyang-Mbo Wildlife Sanctuary, south-western Cameroon. Journal of Tropical Ecology. 229-237. |
| 268 | Niederhauser, C. (2015) All frugivores are not equal: exploitation competition determines seed survival and germination in a fleshy-fruited forest herb. Plant Ecology. 1203-1211. |
| 269 | Noble, J. (1975) The effects of emus (Dromaius novaehollandiae Latham) on the distribution of nitre bush (Nitraria billardieri DC.). Journal of Ecology. |
| 270 | Nogales, M. (1995) Frugivory by Plocama pendula (Rubiaceae) by the rabbit (Oryctolagus cuniculus) in xerophytic zones of Tenerife (Canary Islands). Acta Oecologica. |
| 271 | Nogales, M. (1998) Shrikes, lizards and Lycium intricatum (Solanaceae) fruits: a case of indirect seed dispersal on an oceanic island (Alegranza, Canary Islands). Journal of Ecology. |
| 272 | Nogales, M. (2001) Ecological and biogeographical implications of Yellow-Legged Gulls (Larus cachinnans Pallas) as seed dispersers of Rubia fruticosa Ait. (Rubiaceae) in the Canary Islands. Journal of Biogeography. 1137-1145. |
| 273 | Nogales, M. (2005) Effect of Native and Alien Vertebrate Frugivores on Seed Viability and Germination Patterns of Rubia fruticosa (Rubiaceae) in the Eastern Canary Islands. Functional Ecology. |
| 274 | Nogales, M. (2006) Native dispersers induce germination asynchrony in a macaronesian endemic plant (Rubia fruticosa, Rubiaceae) in xeric environments of the Canary Islands. Journal of Arid Environments. 357-363. |
| 275 | Nowak, J. (2012) It is Good to be Eaten by a Bear: Effects of Ingestion on Seed Germination. The American Midland Naturalist. |
| 276 | Orrock, J. (2005) The effect of gut passage by two species of avian frugivore on seeds of pokeweed, Phytolacca americana. Canadian Journal of Botany. 427-431. |

| | |
|---|---|
| 277 | Otani, T. (2000) Seed dispersal and predation by Yakushima macaques, Macaca fuscata yakui, in a warm temperate forest of Yakushima Island, southern Japan. Ecological Research. 133-144. |
| 278 | Otani, T. (2004) Effects of macaque ingestion on seed destruction and germination of a fleshy-fruited tree, Eurya emarginata. Ecological Research. 495-501. |
| 279 | PadrvÉ¬≥n,B. (2011) Integration of invasive Opuntia spp. by native and alien seed dispersers in the Mediterranean area and the Canary Islands. Biological invasions. 831-844. |
| 280 | Panetta, F. (1997) Recruitment of the invasive ornamental, Schinus terebinthifolius, is dependent upon frugivores. Austral Ecology. 432-438. |
| 281 | Paulsen, T. (2002) Passage through bird guts increases germination rate and seedling growth in Sorbus aucuparia. Functional Ecology. 608-616. |
| 282 | Pegg, N. (2014) Antelope ingestion enhances germination of the marula (Sclerocarya birrea), an important African savannah tree. African Journal of Ecology. 499-505. |
| 283 | Pereira, M. (2011) Effects of bird ingestion on seed germination of Vaccinium cylindraceum(Ericaceae), an endemic species of the Azores archipelago. Botany. |
| 284 | Petre, C. (2015) Western lowland gorilla seed dispersal: Are seeds adapted to long gut retention times?. Acta Oecologica. 59-65. |
| 285 | Pratolongo, P. (2003) Comparative analysis of variables associated with germination and seedling establishment for Prosopis nigra (Griseb.) Hieron and Acacia caven (Mol.) Mol.. Forest Ecology and Management. 15-25. |
| 286 | RamvÉ¬≠rez, M. (2012) Cross-Infection Experiments of Psittacanthus schiedeanus: Effects of Host Provenance, Gut Passage, and Host Fate on Mistletoe Seedling Survival. Plant Disease. |
| 287 | Ramires, M. (2009) Germination of Psittacanthus schiedeanus (mistletoe) seeds after passage through the gut of Cedar Waxwings and Grey Silky-flycatchers1. The Journal of the Torrey Botanical Society. |
| 288 | Reid, S. (2011) Avian gut-passage effects on seed germination of shrubland species in Mediterranean central Chile. Plant Ecology. 1-10. |
| 289 | Renison, D. (2009) The effect of passage through the gut of the Greater Rhea (Rhea americana) on germination of tree seeds: implications for forest restoration. Emu. |
| 290 | Rick, C. (1961) Galapagos tomatoes and tortoises. Evolution. |
| 291 | Righini, N. (2004) Effect of different primate species on germination of Ficus (Urostigma) seeds. Zoo Biology. |
| 292 | Robinson, W. (1986) Effect of Fruit Ingestion on Amelanchier seed Germination. Bulletin of the Torrey Botanical Club. |
| 293 | Rodrigues-Perez, J. (2005) Effect of seed passage through birds and lizards on emergence rate of mediterranean species: differences between natural and controlled conditions. Functional Ecology. |
| 294 | Roehm, K. (2013) Is the Coyote (Canis latrans) a Potential Seed Disperser for the American Persimmon (Diospyros virginiana)?. The American Midland Naturalist. |
| 295 | Rojas-Martinez, A. (2015) Effects of seed ingestion by the lesser long-nosed bat Leptonycteris yerbabuenae on the germination of the giant cactus Isolatocereus dumortieri. The Southwestern Naturalist. 85-89. |
| 296 | Rosalino, L. (2010) The Role of Carnivores as Mediterranean Seed Dispersers. Annales Zoologici Fennici. 47. 195-205. |
| 297 | Rost, J. (2012) Seed dispersal by carnivorous mammals into burnt forests: An opportunity for non-indigenous and cultivated plant species. Basic and Applied Ecology. 623-630. |
| 298 | Rust, R. (1981) Seed production and seedling establishment in the Mayapple, Podophyllum peltatum L.. American Midland Naturalist. |
| 299 | Sato, H. (2012) Frugivory and seed dispersal by brown lemurs in a Malagasy tropical dry forest. Biotropica. |
| 300 | Schaumann, F. (2002) Endozoochorous seed dispersal by martens (Martes foina, M. martes) in two woodland habitats. Flora. 370-378. |
| 301 | Schetini de Azevedo, C. (2013) Effect of passage through the gut of Greater Rheas on the germination of seeds of plants of cerrado and caatinga grasslands. EMU. 177-182. |
| 302 | Setlalekgomo, M. (2014) Seed dispersal by serrated tortoises (Psammobates Oculiferus) and the effect of their gut passage on seed germination. Scientific Journal of Animal Science. |
| 303 | Setlalekgomoa, M. (2013) Effects of gut passage by kudu (Tragelaphus strepsiceros) and pericarp on the germination percentage of morula (Sclerocarya birrea) seeds. Scientific Journal of Biological Sciences. |

| | |
|---|---|
| 304 | Shanahan (2000) Fig-Eating by Bornean Tree Shrews (Tupaia spp.): Evidence for a Role as Seed Dispersers1. Biotropica. |
| 305 | Silverstein, R. (2005) GERMINATION OF NATIVE AND EXOTIC PLANT SEEDS DISPERSED BY COYOTES (CANIS LATRANS) IN SOUTHERN CALIFORNIA. The Southwestern Naturalist. |
| 306 | Soto-Gamboa (2002) Fruit-disperser interaction in a mistletoe-bird system: a comparison of two mechanisms of fruits processing on seed germination. Plant Ecology. |
| 307 | Stevenson, Pablo (2000) Seed dispersal by woolly monkeys (Lagothrix lagothricha) at Tinigua National Park, Colombia: Dispersal distance, germination rates, and dispersal quantity. American Journal of Primatology. |
| 308 | Stevenson, Pablo (2002) Effects of Seed Dispersal by Three Ateline Monkey Species on Seed Germination at Tinigua National Park, Colombia. International Journal of Primatology. |
| 309 | Takasaki, H. (1983) Seed dispersal by chimpanzees: a preliminary note.. African Study Monographs. 3. 105-108. |
| 310 | Tassin, J. (2008) Effect of Ingestion by Drepanoptila holosericea (Columbidae) on the Seed Germination of Santalum austrocaledonicum (Santalaceae). Journal of Tropical Ecology. |
| 311 | Tassin,J. (2010) Can Ptilinopus greyii (Columbidae) Disperse Seeds in New Caledonia's Dry Forests?. Pacific Science. |
| 312 | Thabethe, V. (2015) Ingestion by an invasive parakeet species reduces germination success of invasive alien plants relative to ingestion by indigenous turaco species in South Africa. Biological invasions. 3029-3039. |
| 313 | Traveset (2008) Seed trait changes in dispersers' guts and consequences for germination and seedling growth. Ecology. 95-106. |
| 314 | Traveset, A. (1997) Effects of birds and bears on seed germination in the temperate rainforests of Southeast Alaska. Oikos. |
| 315 | Traveset, A. (1998) Ecology of the fruit-colour polymorphism in Rubus spectabilis. Evolutionary Ecology. 331-345. |
| 316 | Traveset, A. (2001) Ecology of fruit-colour polymorphism in Myrtus communis and differential effects of birds and mammals on seed germination and seedling growth. Journal of Ecology. 749-760. |
| 317 | Traveset, A. (2001) Passage through bird guts causes interspecific differences in seed germination characteristics. Functional Ecology. 669-675. |
| 318 | Valenta, K. (2009) Effects of gut passage, feces, and seed handling on latency and rate of germination in seeds consumed by capuchins (Cebus capucinus). American Journal of Primatology. 486-492. |
| 319 | Valido, A. (1994) Frugivory and seed dispersal by the lizard Gallotia galloti (Lacertidae) in a xerix habitat of the Canary Islands. Oikos. |
| 320 | Varela, O. (2002) Seed Dispersal by Chelonoidis chilensis in the Chaco Dry Woodland of Argentina. Journal of Herpetology. |
| 321 | Varela, O. (2006) Passage time, viability, and germination of seeds ingested by foxes. Journal of Arid Environments. 67. 566-578. |
| 322 | Vila, M. (1998) FRUIT CHOICE AND SEED DISPERSAL OF INVASIVE VS. NONINVASIVE CARPOBROTUS (AIZOACEAE) IN COASTAL CALIFORNIA. Ecology. |
| 323 | Voigt, F.A. (2011) Interactions between the invasive tree Melia azedarach (Meliaceae) and native frugivores in South Africa. Journal of Tropical Ecology. 355-363. |
| 324 | Waibel, A. (2013) Does a giant tortoise taxon substitute enhance seed germination of exotic fleshy-fruited plants?. Journal of Plant Ecology. 57-63. |
| 325 | Walker, L. (1990) Germination of an Invading Tree Species (Myrica faya) in Hawaii. Biotropica. |
| 326 | Webber, B. (2004) Cassowary frugivory, seed defleshing and fruit fly infestation influence the transition from seed to seedling in the rare Australian rainforest tree, Ryparosa sp. nov. 1 (Achariaceae). Functional Plant Biology. 505-516. |
| 327 | Westcott, D. (2007) Cassowary dispersal of the invasive pond apple in a tropical rainforest: the contribution of subordinate dispersal modes in invasion. Diverstiy and Distribution. |
| 328 | White, E. (2006) Murraya paniculata: what is the potential for this popular ornamental plant to become an environmental weed?. Australian Weeds Conference. |
| 329 | Whitney, K. (1998) Seed dispersal by Ceratogymna hornbills in the Dja Reserve, Cameroon. Journal of Tropical Ecology. 351-371. |

| | |
|---|---|
| 330 | Williams, P. (2000) Small mammals as potential seed dispersers in New Zealand. Austral Ecology. 523-532. |
| 331 | Willson, M. (1996) Seed dispersal by lizards in Chilean rainforest. Revista Chilena de Historia Natural. |
| 332 | Wilson, A. (2012) Knysna Turacos (Tauraco corythaix) do not improve seed germination of ingested fruit of some indigenous South African tree species. South African Journal of Botany. 55-62. |
| 333 | Wotton, D. (2002) Effectiveness of the common gecko (Hoplodactylus maculatus) as a seed disperser on Mana Island, New Zealand. New Zealand Journal of Botany. 639-647. |
| 334 | Wrangham, R. (1994) Seed dispersal by forest chimpanzees in Uganda. Journal of Tropical Ecology. 355-368. |
| 335 | Yagihashi, T. (1998) Effects of Bird Ingestion on Seed Germination of Sorbus commixta. Oecologia. |
| 336 | Yagihashi, T. (1999) Effects of bird ingestion on seed germination of two Prunus species with different fruit-ripening seasons. Ecological Research. 71-76. |
| 337 | Zhan-Hui, T. (2007) Effect of Ingestion by Two Frugivorous Bat Species on the Seed Germination of Ficus racemosa and F. hispida (Moraceae). Journal of Tropical Ecology. |
| 338 | Zhan-Hui, T. (2008) Seed Dispersal of Morus macroura (Moraceae) by Two Frugivorous Bats in Xishuangbanna, SW China. Biotropica. |
| 339 | Zhou, Y. (2008) Frugivory and seed dispersal by a small carnivore, the Chinese ferret-badger, Melogale moschata, in a fragmented subtropical forest of central China. Forest Ecology and Management. 1595-1603. |

**Table S2.** Full model results.

| | Coefficient | Estimate | Confidence Interval |
|---|---|---|---|
| Intercept | | 0.2 | (-0.08 - 0.49) |
| Control type | whole fruit | 1.06 | (1.01 - 1.11) |
| Feeding trial | field-collected | 0.06 | (-0.06 - 0.18) |
| | greenhouse soil | -0.3 | (-0.4 - -0.19) |
| | field soil | -0.32 | (-0.43 - -0.22) |
| Sowing medium | other | 0.2 | (-0.13 - 0.53) |
| | other mammals | -0.61 | (-0.98 - -0.24) |
| | primates | 0.89 | (0.42 - 1.37) |
| | reptiles | -0.2 | (-0.48 - 0.07) |
| | bats | 0.41 | (-0.14 - 0.96) |
| | invertebrates | -2.61 | (-3.98 - -1.23) |
| Frugivore taxon | fish | 0.39 | (-0.95 - 1.73) |
| Frugivore invasive? | yes | 0.04 | (-0.49 - 0.57) |
| Plant invasive? | yes | 0.11 | (-0.46 - 0.68) |
| | temperate | 0.47 | (0.24 - 0.69) |
| Latitude region | subtropical | 0.54 | (0.33 - 0.74) |
| Mainland vs. island | island | -0.49 | (-0.64 - -0.35) |

**Table S3.** Full phylogenetic model results.

| | Coefficient | Estimate | Confidence Interval |
|---|---|---|---|
| Intercept | | 0.03 | (-4.87 - 4.92) |
| Control type | whole fruit | 1.06 | (1.01 - 1.11) |
| Feeding trial | field-collected | 0.1 | (-0.02 - 0.22) |
| | greenhouse soil | -0.32 | (-0.43 - -0.21) |
| | field soil | -0.31 | (-0.42 - -0.21) |
| Sowing medium | other | 0.19 | (-0.14 - 0.52) |
| | other mammals | -0.53 | (-0.91 - -0.16) |
| | primates | 0.67 | (0.18 - 1.15) |
| | reptiles | -0.2 | (-0.47 - 0.08) |
| | bats | 0.25 | (-0.3 - 0.81) |
| | invertebrates | -2.32 | (-3.91 - -0.73) |
| Frugivore taxon | fish | 0.25 | (-1.14 - 1.63) |
| Frugivore invasive? | yes | 0 | (-0.54 - 0.54) |
| Plant invasive? | yes | 0.22 | (-0.29 - 0.73) |
| | temperate | 0.49 | (0.26 - 0.72) |
| Latitude region | subtropical | 0.59 | (0.38 - 0.8) |
| Mainland vs. island | island | -0.44 | (-0.58 - -0.3) |

Table S4. Results of linear hypothesis tests.

| Test | Estimate | Std. Error | z value | P value |
|---|---|---|---|---|
| *Sowing medium* | | | | |
| Petri - Field | 0.323 | 0.054 | 5.979 | <0.001 |
| Petri - Nursery | 0.295 | 0.055 | 5.325 | <0.001 |
| Petri - Other | -0.198 | 0.168 | -1.175 | 0.24 |
| Field - Nursery | -0.027 | 0.057 | -0.476 | 0.634 |
| Field - Other | -0.52 | 0.17 | -3.057 | 0.002 |
| Nursery - Other | -0.493 | 0.161 | -3.053 | 0.002 |
| *Feeding trial* | | | | |
| Feeding trial - Field collection | 0.06 | 0.061 | 0.99 | 0.322 |
| *Scarification vs. de-inhibition* | | | | |
| Scarification - De-inhibition | -1.059 | 0.025 | -41.756 | <0.001 |
| *Frugivore taxon* | | | | |
| Bird - Bat | -0.407 | 0.28 | -1.452 | 0.147 |
| Bird - Primate | -0.889 | 0.242 | -3.679 | <0.001 |
| Bird - Other mammal | 0.607 | 0.185 | 3.281 | 0.001 |
| Bird - Reptile | 0.202 | 0.141 | 1.438 | 0.15 |
| Bird - Fish | -0.404 | 0.679 | -0.595 | 0.552 |
| Bird - Invertebrate | 2.602 | 0.703 | 3.702 | <0.001 |
| Bat - Primate | -0.482 | 0.334 | -1.446 | 0.148 |
| Bat - Other | 1.014 | 0.302 | 3.357 | 0.001 |
| Bat - Reptile | 0.609 | 0.304 | 2.005 | 0.045 |
| Bat - Fish | 0.003 | 0.722 | 0.004 | 0.997 |
| Bat - Invertebrate | 3.009 | 0.745 | 4.041 | <0.001 |
| Primate - Other | 1.496 | 0.264 | 5.666 | <0.001 |
| Primate - Reptile | 1.091 | 0.267 | 4.081 | <0.001 |
| Primate - Fish | 0.485 | 0.703 | 0.69 | 0.49 |
| Primate - Invertebrate | 3.491 | 0.729 | 4.786 | <0.001 |
| Other - Reptile | -0.405 | 0.218 | -1.853 | 0.064 |
| Other - Fish | -1.011 | 0.691 | -1.462 | 0.144 |
| Other - Invertebrate | 1.995 | 0.71 | 2.808 | 0.005 |
| Reptile - Fish | -0.606 | 0.689 | -0.879 | 0.379 |
| Reptile - Invertebrate | 2.399 | 0.713 | 3.368 | 0.001 |
| Fish - Invertebrate | 3.006 | 0.967 | 3.108 | 0.002 |
| *Latitude region* | | | | |
| Tropical - Subtropical | -0.537 | 0.104 | -5.157 | <0.001 |
| Tropical - Temperate | -0.468 | 0.113 | -4.128 | <0.001 |
| Subtropical - Temperate | 0.069 | 0.069 | 1.004 | 0.315 |
| *Mainland vs. island* | | | | |
| Mainland - Island | 0.493 | 0.074 | 6.702 | <0.001 |
